# Dynamical modelling of the heat shock response in *Chlamydomonas reinhardtii*

**DOI:** 10.1101/085555

**Authors:** Stefano Magni, Antonella Succurro, Alexander Skupin, Oliver Ebenhöh

**Affiliations:** Institute for Quantitative and Theoretical Biology, Heinrich Heine University Düsseldorf, Germany; Luxembourg Centre for Systems Biomedicine, University of Luxembourg, Esch-sur-Alzette, Luxembourg; Cluster of Excellence on Plant Sciences (CEPLAS), Cologne Biocenter, University of Cologne, Germany; Cluster of Excellence on Plant Sciences (CEPLAS), Institute for Quantitative and Theoretical Biology, Heinrich Heine University Düsseldorf, Germany

**Keywords:** heat shock response, dynamical modelling, Chlamydomonas reinhardtii, heat shock proteins

## Abstract

Global warming is exposing plants to more frequent heat stress, with consequent crop yield reduction. Organisms exposed to large temperature increases protect themselves typically with a heat shock response (HSR). To study the HSR in photosynthetic organisms we present here a data driven mathematical model describing the dynamics of the HSR in the model organism *Chlamydomonas reinhartii*. Temperature variations are sensed by the accumulation of unfolded proteins, which activates the synthesis of heat shock proteins (HSP) mediated by the heat shock transcription factor HSF1. Our dynamical model employs a system of ordinary differential equations mostly based on mass-action kinetics to study the time evolution of the involved species. The signalling network is inferred from data in the literature, and the multiple experimental data-sets available are used to calibrate the model, which allows to reproduce their qualitative behaviour. With this model we show the ability of the system to adapt to temperatures higher than usual during heat shocks longer than three hours by shifting to a new steady state. We study how the steady state concentrations depend on the temperature at which the steady state is reached. We systematically investigate how the accumulation of HSPs depends on the combination of temperature and duration of the heat shock. We finally investigate the system response to a smooth variation in temperature simulating a hot day.

## 1 Introduction

As a consequence of global warming, plants are more and more subject to heat stress, a condition that can severely reduce crop yield [Lobell et al., 2011, Deryng et al., 2014]. Understanding how plants react to such a stress is of crucial importance in developing metabolic engineering approaches or treatments to improve crop plant resistance to heat. In general when exposed to increased temperature organisms react with a heat shock response (HSR), which to a certain degree allows adaptation to the new conditions. The proteins involved in the HSR are highly conserved among organisms ranging from bacteria to mammals [Boorstein et al., 1994, Gupta, 1995]. Moreover, because temperature affects the cell as a whole, HSRs cover a wide range of cellular activities localised in all intracellular compartments [Verghesea et al., 2012, Velichko et al., 2013]. Depending on organism and cell type, HSRs are activated by a number of different sensing mechanisms [Richter et al., 2010].

The green microalgae *Chlamydomonas reinhardtii* is a widely studied, easy to grow photosynthetic model organism, for which techniques have been established to modify all three (nuclear, mitochondrial and plastidial) genomes [Rochaix, 1995, Duby and Matagne, 1999]. It thus represents an ideal candidate to study the mechanisms involved in the HSR of plants. In addition, *C. reinhardtii* is also interesting on its own for possible industrial applications, such as the production of biopharmaceuticals, biofuels and hydrogen. Different aspects of the HSRs in *C. reinhardtii* have been experimentally investigated, as reviewed e.g. in [Schroda et al., 2015].

Both in the land plant *Arabidopsis thaliana* [Kurepa et al., 2003, Sugio et al., 2009] and in the green alga *C. reinhardtii* [Schmollinger et al., 2013] the heat shock response is generally triggered by a heat-induced accumulation of mis-or unfolded proteins and leads, through a series of sensor and signalling events, to the activation of a heat shock transcription factor (HSF). The HSF in turn promotes the expression of heat shock protein (HSP) genes, and subsequently of the synthesis of proteins, some of which act as chaperones responsible to refold the degenerated proteins back to their correct three-dimensional structure [Craig et al., 1993]. The precise temperature at which the denatured proteins accumulate depends on the typical temperature range in which an organisms grows [Lindquist and Craig, 1988]. For *C. reinhardtii* it has been shown that a temperature of *T*_0_ = 36°C is sufficient to detect a heat shock response [Kobayashi et al., 2014].

The considerable experimental effort performed until now to study the HSR is in stark contrast to the rather limited complementary theoretical activities aiming at developing mathematical models of the HSR. However, mathematical models are increasingly recognised as a powerful tool to investigate dynamic biological systems (as reviewed e.g. in [Pfau et al., 2011]). The construction of a mathematical model itself often provides a high degree of insight, because it forces the researchers to focus on the essential features of the system under investigation and thus to identify the key components which are responsible for the generation of characteristic system properties. It thus allows to discriminate between important and less important and provides a powerful technique to verify whether our general understanding of a system is basically correct and whether the interacting molecular mechanisms that have been identified experimentally are sufficient to reproduce and thus explain observed responses. The importance of mathematical modelling in understanding and unveiling the functioning of plant signalling processes has recently been highlighted in [Chew et al., 2014].

One of the earliest theoretical studies of the eukaryotic HSR took mainly the influence of mis-folded proteins into account and did not include a detailed transcriptional regulation [Peper et al., 1998]. Modelling of the transcriptional regulation was firstly used to study procaryotic systems, in particular *E. coli*, where the transcription factor playing the role of HSF is called *σ*^32^. This was done in [Srivastava et al., 2001] employing a stochastic approach, and in [Kurata et al., 2001] with a deterministic one. More recently, [Rieger et al., 2005] proposed a mathematical model to describe the HSR in HeLa cells with a detailed model for nuclear events. While that model is highly useful to gain a principle understanding of the interacting molecular mechanisms, it uses dimensionless variables, where the dynamics are normalised by the maximal response, which makes possible only relative predictions, and not absolute ones. A model of the thermal adaptation in *Candida albicans*, a fungal pathogen of humans, is presented in [Leach et al., 2012b]. That model mainly focuses on the auto-regulatory mechanism involving HSF1 and HSP90 (see next section), but it does not include a detailed transcriptional regulation. The role of HSP90 and its interactions with HSF1 are further described in [Leach et al., 2012a]. The modelling of the multi-scale heat stress response in the budding yeast *Saccharomyces cerevisiae* is discussed in [Fonseca et al., 2012]. Recently, [Sivéry et al., 2015] focused on the role of HSF1 during the HSR in mammals.

As argued in [Matuszyńska and Ebenhöh, 2015], every mathematical model is usually constructed with a particular purpose and research questions it should help to answer. A model serves as an *in silico* workbench that can be used to simulate the behaviour of the modelled system in very diverse situations. This allows exploring a variety of scenarios potentially difficult to test, or not yet tested, experimentally. Such simulations often allow the proposition of new hypothesis on the functioning of the biological system under investigation, and to suggest which new experiments could be performed to test these hypotheses. Thus, models provide a theoretical framework to complement the experimental effort.

In this work we build a data driven mathematical model for the HSR in the green algae *Chlamydomonas reinhartii* with the main purpose to i) verify whether our understanding of the mechanisms of HSR are not only qualitatively but also quantitatively (in relative, not absolute, terms) consistent with experimental observations, and to ii) provide a generic theoretical framework, by which new predictions can be made (such as responses to chemical treatments or genetic modifications) and thus novel hypotheses can be generated. In Section 2 we introduce the signalling network used to implement the mathematical description of the HSR and we discuss the typical behaviour of the model. In Section 3 we compare simulation results with experimental data from the literature, focussing on experiments in which specific inhibitors were applied and on “double heat shock” experiments, in which a second heat shock was applied after a certain relaxation period. We discuss in particular how the results are used for model calibration. In Section 4 we employ the calibrated model to simulate interesting conditions that have not yet been tested experimentally, and we thus demonstrate the predictive power of the model and its usefulness in providing a fundamental understanding of the systems dynamics and its emergent properties. In Section 5 we discuss how our model could possibly be extended to include the activation of the HSR by a shift from dark to light of the cells, and we conclude.

## 2 Model development

Heat shock proteins are molecular chaperones transiently expressed in response to heat stress to maintain protein homeostasis, and most families are conserved among different species [Richter et al., 2010]. There exist a number of different heat shock protein families, which are typically named HSP*w* with the integer number *w* indicating the molecular weight of the protein. *C. reinhardtii* can synthesize different families of heat shock proteins (see *e.g*. [Schroda, 2004]), of which the families HSP70 and HSP90 are of the main interest in the current work. The HSP70 family comprises cytoplasmic HSP70A [Müller et al., 1992], plastidic HSP70B [Drzymalla et al., 1996] and mitochondrial HSP70C, while the HSP90 family comprises cytosolic HSP90A, HSP90B in the Endoplasmic Reticulum and HSP90C in the chloroplast [Willmund and Schroda, 2005].

We develop our mathematical model with the goal to provide a generic description of the heat shock response. Therefore, we decide to simplify the model by including only one generic heat shock protein (indicated by HP), which can represent any HSP present in *C. reinhardtii*. The simulation results are consequently compared with data relative to various of the above mentioned HSPs. In *C. reinhardtii* the only HSF (among the two encoded in the genome) known to be activated by heat shock is HSF1, characterized in [Schulz-Raffelt et al., 2007]. Therefore, HSF1 corresponds to the HSF described in our model. In land plants at least 18 different HSF are present [Scharf et al., 2012], which is an example of the fact that in general gene families in *C. reinhardtii* are smaller than in land plants (see for instance [Schroda, 2004] for another such example concerning chaperones). This simplicity further supports the choice of *C. reinhardtii* as a suitable model organism to study the heat shock response in plants.

### 2.1 Modelling the signalling network

From the experiments performed in [Schmollinger et al., 2013] we derive the signalling network schematically depicted in Fig. 1. All components are described in detail in Table 1. In [Schmollinger et al., 2013] it has been shown that the temperature increase triggering the HSR is sensed by the accumulation of degenerated proteins P#. Their presence activates a stress kinases SK, which, in the active form SK*, phosphorylates the heat shock factor HSF. The phosphorylated (HSF*) and un-phosphorilated (HSF) heat shock factor can bind to the transcription factor binding sites of various genes, coding for key proteins involved in the HSR, including HSF itself and heat shock protein (HP). In the model, all these genes are described by one variable G, and the transcription of different mRNAs is represented by the individual transcription rates *π*. Binding of the active form HSF* to genes G induces the production of mRNA coding for heat shock protein (mR_HP_) and for the heat shock factor itself (mR_F_), whereas the inactive form HSF blocks the transcription. The mRNAs are subsequently translated into the corresponding proteins HP and HSF (with rates *η*), respectively. The increase in HSF concentration leads to a higher occupation of the gene with the inactive form. The increased concentration of chaperones HP increases the repair of the degenerated protein state P# to their functional form P until a new steady state is reached.

**Figure 1:**
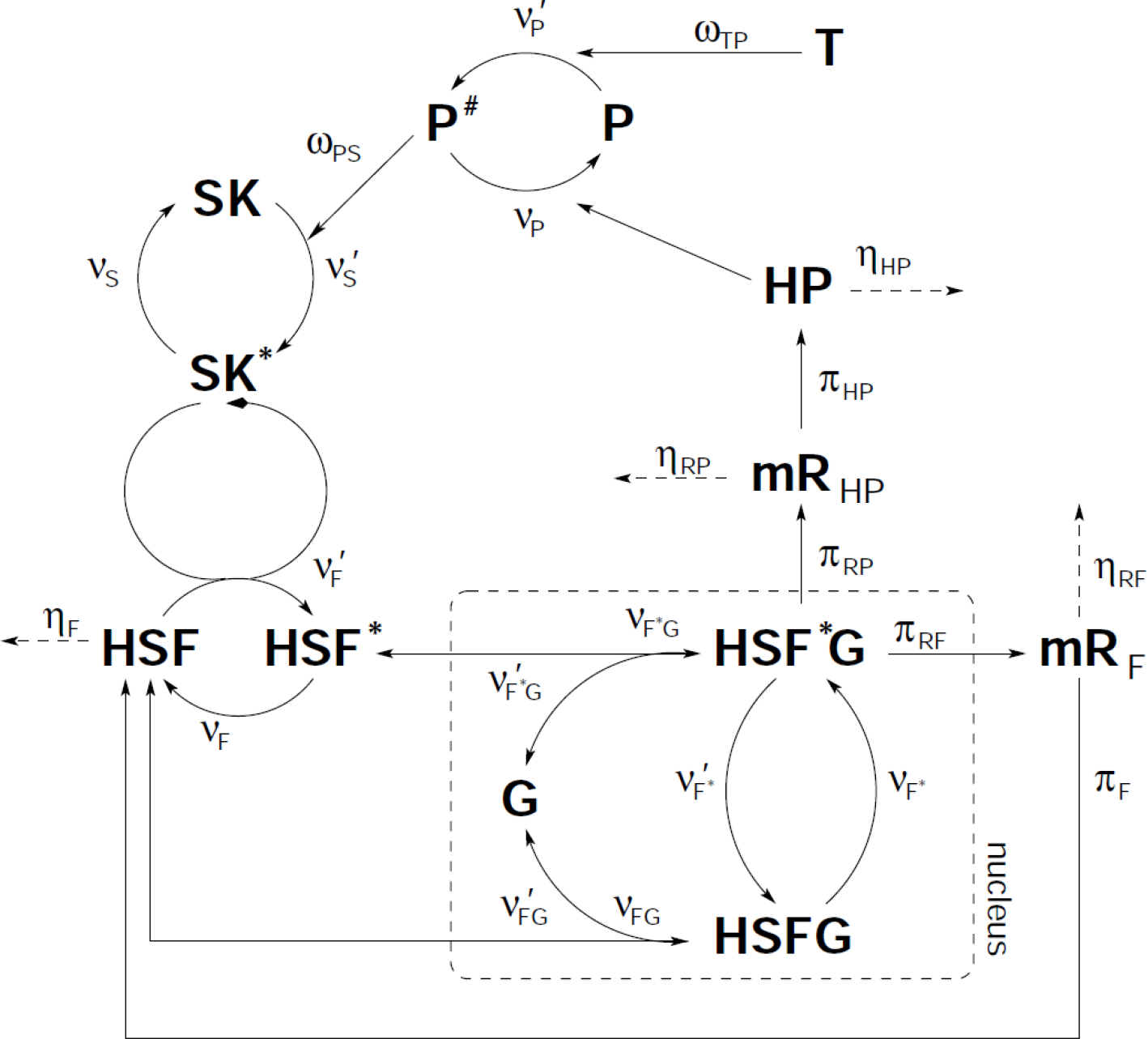
Scheme of the signalling network that we use to model the HSR. Temperature T acts via the Arrhenius law *ω*_TP_ on the protein level P. Higher temperature increases *ν*′_P_ leading to more degenerated proteins P#. This activates stress kinases SK by a hill kinetics *ω*_PS_ which increases phosphorilization of the heat shock factor HSF. If HSF is bound to the gene G, mRNA for the heat shock factor HSF and for the heat shock protein HP is generated by the corresponding production rates *π*. The mRNA is translated into the proteins HP and HSF and degraded by rates *η*.

**Table 1:** Variables and names used in the model.

| Variable | Representing<br>concentration of |
| --- | --- |
| $[P]$ | Protein |
| $[P^\#]$ | Degenerated protein |
| $[SK]$ | Stress kinases (SK) |
| $[SK^*]$ | Phosphorylated SK |
| $[HSF]$ | Heat shock factor (HSF) |
| $[HSF^*]$ | Phosphorylated HSF |
| $[G]$ | Free gene |
| $[HSF^*G]$ | HSF bound to gene, active |
| $[HSFG]$ | HSF bound to gene, inactive |
| $[mR_F]$ | mRNA for the HSF |
| $[mR_{HP}]$ | mRNA for the HSP |
| $[HP]$ | Heat shock protein (HSP) |

**Table 2:** Values of the rate constants used in the model. The values employed in all the simulations shown in this work are those labelled as final values, while those labelled as fiducial values are those employed as a starting point for the optimization procedure described in Section 3.3.

| Rate constant | Fiducial value | Final value | Unit of measure |
| --- | --- | --- | --- |
| $k_P$ | 10 | 11.49 | $(\mu\text{M s})^{-1}$ |
| $k'_P$ | 100 | 111.9 | $\text{s}^{-1}$ |
| $k_S$ | 100 | 100.7 | $\text{s}^{-1}$ |
| $k'_S$ | 500 | 533.1 | $\text{s}^{-1}$ |
| $k'_F$ | 1 | 0.9557 | $\text{s}^{-1}$ |
| $k_F$ | 1 | 0.9640 | $(\mu\text{M s})^{-1}$ |
| $k'_{FG}$ | 0.10 | 0.08371 | $\text{s}^{-1}$ |
| $k_{FG}$ | 0.0050 | 0.005574 | $(\mu\text{M s})^{-1}$ |
| $k_{F^*G}$ | 1 | 0.9131 | $(\mu\text{M s})^{-1}$ |
| $k'_{F^*G}$ | 0.50 | 0.4136 | $\text{s}^{-1}$ |
| $k'_{F^*}$ | 0.010 | 0.01064 | $\text{s}^{-1}$ |
| $k_{F^*}$ | 0.010 | 0.01192 | $\text{s}^{-1}$ |
| $k_{\pi_{RF}}$ | 16 | 18.16 | $\text{s}^{-1}$ |
| $k_{\pi_{RH}}$ | 4.5 | 4.193 | $\text{s}^{-1}$ |
| $k_{\pi_{HP}}$ | 0.5 | 0.5142 | $\text{s}^{-1}$ |
| $k_{\pi_F}$ | 0.02 | 0.02112 | $\text{s}^{-1}$ |
| $d_F$ | 0.001 | 0.0008728 | $\text{s}^{-1}$ |
| $d_{HP}$ | 0.000086 | 0.0009384 | $\text{s}^{-1}$ |
| $d_{RF}$ | 0.0015 | 0.001719 | $\text{s}^{-1}$ |
| $d_{RP}$ | 0.0012 | 0.001017 | $\text{s}^{-1}$ |

The model is described by twelve dynamic variables (see Table 1), each representing the concentration of the corresponding component of the network. The concentrations are expressed in the figures in arbitrary units (a.u.), which are used to normalize each panel to a reference value. These reference values are the same across different figures, and are listed in table 5. They are necessary because, since no data for the absolute values of the concentration of any of the species involved are available, the model variable could be fit only to relative data.

The temporal dynamics of the variables are governed by a set of ordinary differential equations (ODEs), which are reported in Table 3 in the Supplementary Material. For a review covering the applicability of ODEs and other types of modelling to biological systems see *e.g*. [Pfau et al., 2011]. The equations we employ depend on rate expressions (see Table 4 of the Supplementary Material) describing the various regulatory processes (*ω*), activation and deactivation steps (*ν*), synthesis rates (*π*) and degradation rates (*η*). For the majority of these rate laws we assume mass action kinetics. We also employ some additional non-linear terms having a behaviour of the type of Michaelis-Menten kinetics or Hill kinetics, listed in Table 4 of the Supplementary Material. The term following a Hill kinetics is the one describing the regulatory process *ω*_*ps*_ involved in the reaction by which the denatured proteins induce the activation (phosphorylation) of the SK. It allows to have a response with the sigmoidal shape able to describe a threshold effect in the activation of this reaction. The action of the phosphorylated SK*, the enzyme phosphorylating HSF, is described by a Michaelis-Menten behaviour, typical of enzymatic kinetics. The effect of the temperature, which denatures (unfolds or misfolds) proteins, is described by means of the Arrhenius law, with an activation energy in the wide range reported in the literature for the activation energies of protein denaturation due to thermal stress (as discussed in [Bischof and He, 2006, He and Bischof, 2003]). More details on these terms can be found in Table 4 of the Supplementary Material.

**Table 3:**
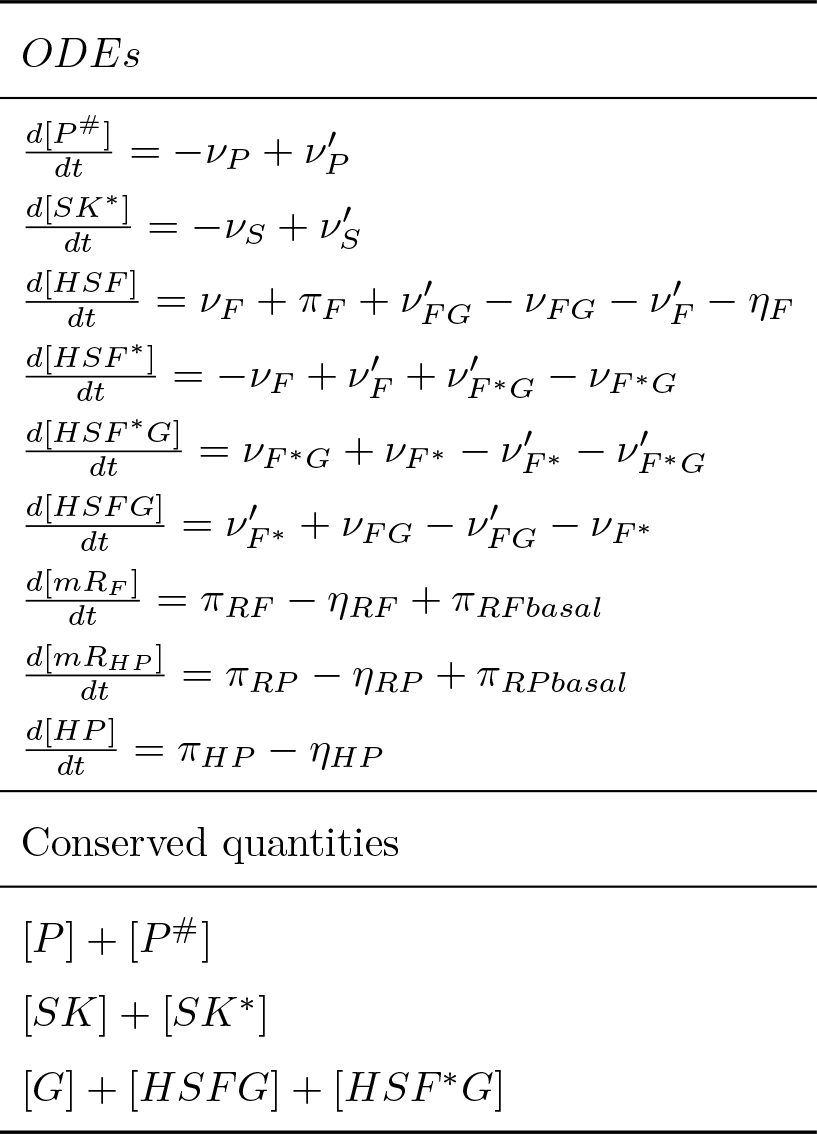
The ODEs used in the model and the conserved quantities. Even if the system has twelve variables (listed in Table 1), only nine ODEs are required to model it. In fact, there are three conserved quantities: [*P*] + [*P*^#^], [*SK*] + [*SK*\*] and [*G*] + [*HSFG*] + [*HSF*G*] are constants. The initial conditions used for the twelve variables are: [*P*] = 100000 *μ*M, [P#] = 1 *μ*M, [*SK*] = 0.1 *μ*M, [*SK\**] = 0.05 *μ*M, [*HSF*] = 10.5 *μ*M, [*HSF*\*] = 1 *μM*, [*G*] = 0.0012 *μ*M, [*HSF*G*] = 0.0002 *μ*M, [*HSFG*] = 0.0008 *μ*M, [*mR*_*F*_] = 0.0036 *μ*M, [*mR*_*HP*_] = 0.0036 *μ*M, [*HP*] = 1 *μ*M. Let us remark that the values of the variables are initiated at the initial conditions above, but before applying any HS we let the system run for a long time, so that it has reached the steady state when we apply any HS. This part of each simulation is not shown in the plots.

**Table 4:** Kinetic rate laws used in the model. The reactions are those represented in the scheme of Fig. 1, and the rate laws are mainly based on mass action kinetics, a part from some terms winch follows Arrhenius law or have a Michaelis-Menten or Hill kinetics behaviour. The last is 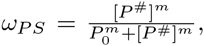, where *m* = 5 and *P*_0_ = 600 *μ*M. The term following the Arrhenius law is 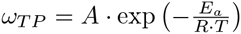, with an activation energy of *E*_*a*_ = 174.440 KJ mol^−1^, perfectly in the wide range reported in the literature for the activation energies of protein denaturation due to thermal stress (as discussed in [Bischof and He, 2006, He and Bischof, 2003]), and with *A* = 9.4318 × 10^28^, *R* = 8.3144598 J mol^−1^ K^−1^ the ideal gas constant and *T* the temperature. Finally, the basal rates are computed as 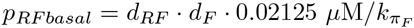 and 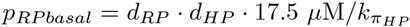.

| Rate law | Reaction |
| --- | --- |
| $\nu_P = k_P \cdot [P^\#] \cdot [HP]$ | $P^\# + HP \rightarrow P + HP$ |
| $\nu'_P = k'_P \cdot [P] \cdot \omega_{TP}$ | $P \rightarrow P^\#$ |
| $\nu_S = k_S \cdot [SK^*]$ | $SK^* \rightarrow SK$ |
| $\nu'_S = k'_S \cdot [SK] \cdot \omega_{PS}$ | $S + P^\# \rightarrow SK^* + P^\#$ |
| $\nu'_F = k'_{HSF} \cdot [HSF] \cdot \frac{[SK^*]}{1+[SK^*]}$ | $HSF + SK^* \rightarrow HSF^* + SK^*$ |
| $\nu_F = k_{HSF} \cdot [HSF^*]$ | $HSF^* \rightarrow HSF$ |
| $\nu'_{HSFG} = k'_{HSFG} \cdot [HSFG]$ | $HSFG \rightarrow HSF + G$ |
| $\nu_{HSFG} = k_{HSFG} \cdot [G] \cdot [HSF]$ | $HSF + G \rightarrow HSFG$ |
| $\nu_{HSF^*G} = k_{HSF^*G} \cdot [G] \cdot [HSF^*]$ | $HSF^* + G \rightarrow HSF^*G$ |
| $\nu'_{F^*G} = k'_{HSF^*G} \cdot [HSF^*G]$ | $HSF^*G \rightarrow HSF^* + G$ |
| $\nu'_{HSF^*} = k'_{HSF^*} \cdot [HSF^*G]$ | $HSF^*G \rightarrow HSFG$ |
| $\nu_{HSF^*} = k_{HSF^*} \cdot [HSFG]$ | $HSFG \rightarrow HSF^*G$ |
| $\pi_{RF} = k_{\pi_{RF}} \cdot [HSF^*G]$ | $HSF^*G : HSF^*G + mRF$ |
| $\pi_{RP} = k_{\pi_{RP}} \cdot [HSF^*G]$ | $HSF^*G : HSF^*G + mRHP$ |
| $\pi_{HP} = k_{\pi_{HP}} \cdot [mR_{HP}]$ | $mR_{HP} : HP + mR_{HP}$ |
| $\pi_F = k_{\pi_F} \cdot [mR_F]$ | $mR_F : HSF + mR_F$ |
| $\eta_F = d_F \cdot [HSF]$ | $HSF \dashrightarrow$ |
| $\eta_{HP} = d_{HP} \cdot [HP]$ | $HP \dashrightarrow$ |
| $\eta_{RF} = d_{RF} \cdot [mR_F]$ | $mR_F \dashrightarrow$ |
| $\eta_{RP} = d_{RP} \cdot [mR_{HP}]$ | $mR_{HP} \dashrightarrow$ |
| $\pi_{RFbasal} = p_{RFbasal}$ | $\rightarrow mR_F$ |
| $\pi_{RPbasal} = p_{RPbasal}$ | $\rightarrow mR_P$ |

Most of the involved rate constants are not known. However, due to the relatively simple model structure, it was rather straight forward to initially choose the rate constants in a biologically reasonable range, and manually fit these parameters to qualitatively fit the experimental data. In a subsequent step, explained in more detail in Section 3.3, this initial manual fit was refined to optimise the reproduction of the data, resulting in the parameter set presented in Table 2 of the Supplementary Material.

### 2.2 Typical behaviour of the model

To investigate the typical behaviour of the model we simulate a heat shock at time *t* = 20 min by suddenly increasing the temperature from 25°C to 42°C This scenario mimics a standard experimental design, in which the temperature is rapidly increased to induce a heat shock response. It should be noted that this is of course an idealised scenario, which can only be approximated experimentally. The time evolution of the concentrations of the molecular species described by the model is depicted in Fig. 2. In this figure, as in the following ones, a red background indicates the period of time during which a heat shocking temperature is provided (42°C in this case). These concentrations are expressed in arbitrary units (a.u.). They are in fact normalized to a standard value for each panel, as reported in table 5 of the Supplementary Material. For the panels A (proteins), B (stress kinases) and D (genes) the normalization factor corresponds to the sum of the amount of the species shown in that plot, which is a conserved quantity for each of these plots.

**Figure 2:**
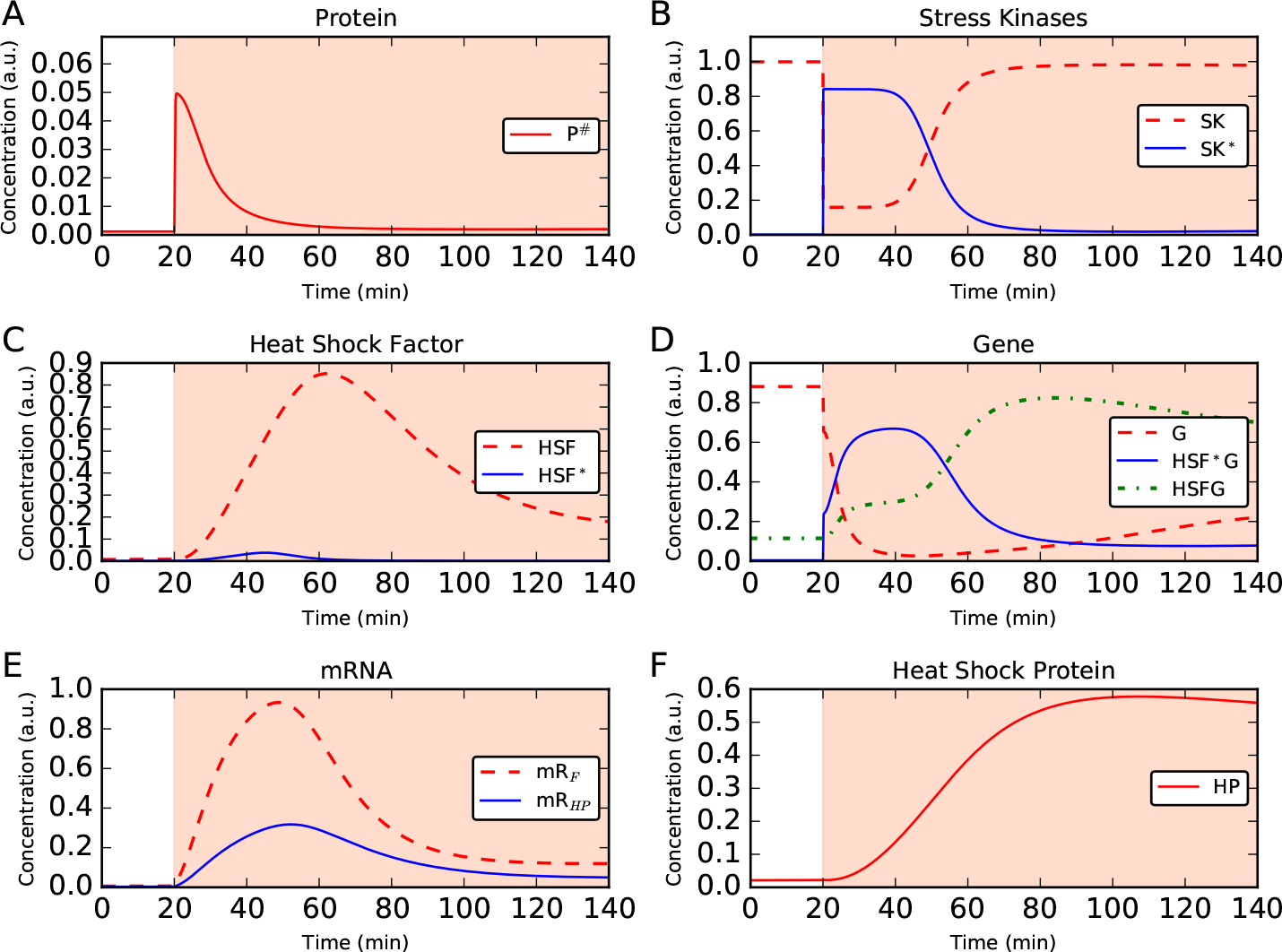
Typical behaviour of the model illustrated inducing a HSR via an increase of the temperature from 25°C to 42°C applied at t = 20 min (represented by a red background in the figures). A: Due to temperature increase at t = 20 min functional proteins P are mis-folded leading to an increased P# level. B: The degenerated proteins bring inactive stress kinases SK into their active form SK*. C: Due to active stress kinases, the heat shock factor (HSF) is phosphorylated (HSF*). D: The heat shock factor HSF binds to free gene loci G, the bound form HSF^*^ G activates mRNA production, and HSF un-binding blocks transcription. E: The initiated gene transcription leads to mRNA production of the HSF and the heat shock protein as shown. F: Due to translation of the corresponding mRNA, the HP concentration increases until the response is switched of. The small degeneration rate of the chaperon leads to a slow decrease after the onset of the HSR. The normalization factors used to represent the concentrations in arbitrary units can be found in table 5 of the Supplementary Material.

**Table 5:**
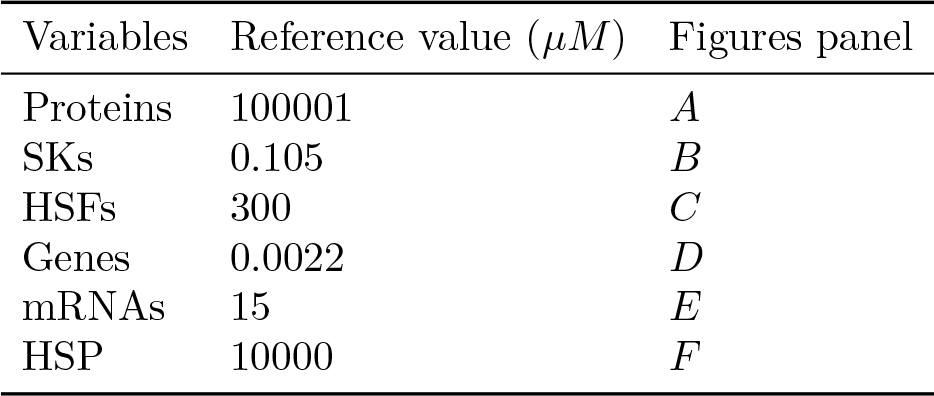
Reference values used to rescale the variables of the model to arbitrary units (a.u.). For Proteins, SKs and Genes the variables are rescaled using the corresponding conserved quantity of Table 3.

Due to the temperature increase, the functional proteins P are mis-folded as shown in panel A. The concentration of the functional form [*P*] of the protein suddenly decreases while that of the mis-folded form [*P*#] increases correspondingly. The latter form induces the transition of the stress kinases SK into their active form SK*, the concentrations of both shown in panel B. Active stress kinases phosphorylate the heat shock factor HSF, shown in panel C. The activated HSF* activates gene transcription as shown in panel D. The amount of free gene G decreases and the active form with phosphorylated HSF bound, HSF*G, increases rapidly. Simultaneously, the gene bound to the inactive form of HSF (HSFG) increases as well, but with a slower dynamics. The activated gene induces mRNA production as shown in panel E for both the mRNAs corresponding to the HSF and to the HSP. These mRNAs are translated, leading to an increase of the heat shock factor itself (panel C) and of the heat shock protein (panel F). The increased chaperon level depicted in panel F leads to re-folding of degenerated proteins into their functional form (as depicted in panel A), which eventually leads to a termination of the response. This analysis illustrates that the model is able to realistically describe the HSR.

## 3 Comparison between simulations and experimental data

In this section we demonstrate that the model is not only able to qualitatively produce a plausible heat shock response, but can also quantitatively reproduce a number of key experimental treatments. We first focus on simulating the feeding experiments performed in [Schmollinger et al., 2013] (Section 3.1), where specific inhibitors have been applied in different concentrations and the effect on the heat shock response was monitored. In Section 3.2 we simulate the double heat shock experiments reported in [Schroda et al., 2000], where a second heat shock was applied after a first initial heat shock to identify the minimum relaxation time needed to observe a full second response. These experiments were used for the fine tuning of the parameters. The calibration procedure is described in Section 3.3.

### 3.1 Inhibitor treatments

In the systematic experiments reported in [Schmollinger et al., 2013], *C. reinhardtii* cells have been fed with different concentrations of specific inhibitors, and the effect on the HSR has been observed by monitoring the temporal evolution of mRNA concentrations of mainly the HSF1 and HSP90 genes. We specifically consider the two inhibitors Staurosporine, a protein kinase inhibitor [Karaman et al., 2008], and Radicicol, a specific inhibitor of HSP90 [Roe et al., 1999]. We simulate these experiments by altering the corresponding rate constants to mimic the effect of the inhibitors, and apply the same heat shock conditions as in the experiments, simulating a sudden temperature increase from 25°C to 40°C at t = 0 min.

#### 3.1.1 Staurosporine

Staurosporine is a protein kinase inhibitor. We therefore simulate the effect of applying Staurosporine by lowering the rate constant *k*′_*F*_, which determines the reaction rate *v*′_*F*_, by which the stress kinase SK activates the HSF. The simulation results are depicted in the left panel of Fig. 3, together with redrawn experimental data from Fig. 1B of [Schmollinger et al., 2013] in the right panel.

**Figure 3:**
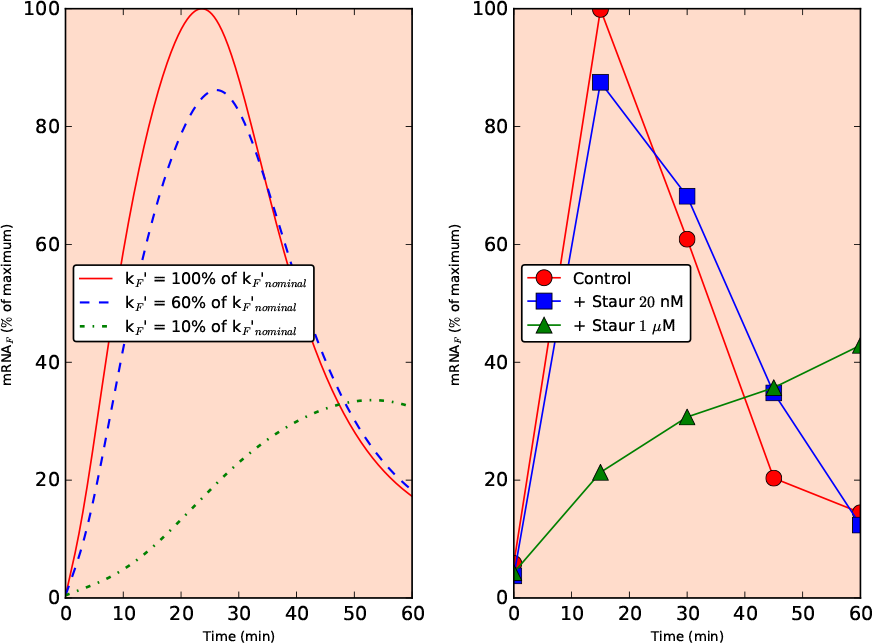
Simulation of the feeding with Staurosporine experiment (left) and comparison with the corresponding data from [Schmollinger et al., 2013] (right).

The simulation results (left panel of Fig. 3) shows the effect of decreasing the rate constant *k*′_*F*_ on the dynamics of the HSF mRNA concentration. Clearly, decreasing the rate constant leads to a reduced maximal mRNA concentration and a delayed response. As can be seen in the right panel of Fig. 3, the same behaviour is observed in the experiments: an increased inhibitor concentration leads to a reduced and delayed transcription of HSF mRNA. Moreover, it can be observed that for a small decrease of the rate constant the response is qualitatively unaltered, whereas a larger decrease corresponding to higher Staurosporine concentration has a qualitative impact, leading to a long delay and considerable reduction of mRNA levels.

Because the experimental data is not quantitative but normalised to the maximal response, a direct comparison of the simulated concentrations is not possible. Moreover, the applied staurosporine concentration can not directly be translated into a reduced rate constant. However, the simulated responses for *k*′_*F*_ at its nominal value of 1.09 *s*^−1^ and for values reduced to 60% and 10% of that value lead to a remarkable agreement between simulation and experiment, in which Staurosporine was applied in concentrations of 20 *n*M and 1 *μ*M, respectively. Not only the qualitative behaviour is well captured, but also the timing of the response as well as the relative reduction of the mRNA signal is quantitatively reproduced.

#### 3.1.2 Radicicol

Radicicol is a specific inhibitor of HSP90 activity. Therefore, we simulate the effect of Radicicol by lowering the rate constant *k*_*P*_, which determines the reaction rate *v*_*P*_, by which in our model the *HSP* refold the unfolded proteins P# back to their original form. The simulation results and the corresponding data (reproduced from Fig. 4B of [Schmollinger et al., 2013]) are displayed in Fig. 4.

**Figure 4:**
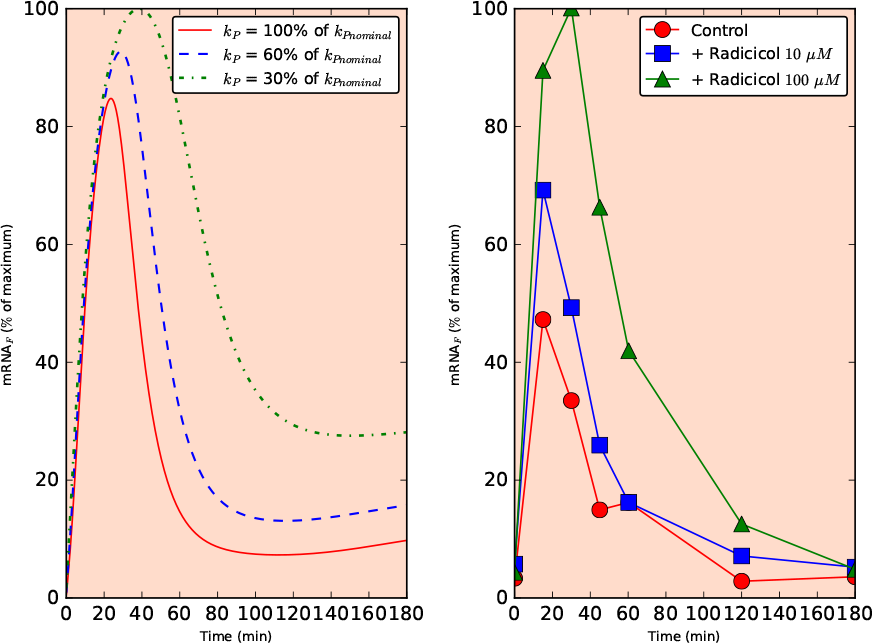
Simulation of the feeding with Radicicol experiment (left) and comparison with the corresponding data from [Schmollinger et al., 2013] (right).

The simulated dynamics of the HSF mRNA concentration is depicted in the left panel of Fig. 4 for three values of the rate constant *k*_*P*_, corresponding to the reference value of 9.938 (*μMs*)^−1^, and for a reduction to 60% and 30% of that value. It can be observed that decreasing the rate constant results to an increased amplitude and delayed attenuation of the HSR. The data from [Schmollinger et al., 2013] shown in the right panel for a control and Radicicol concentrations of 10 *μM* and 100 *μM* demonstrates that a similar behaviour is observed in the experiments. Interestingly, the magnitude of the responses are only qualitatively reproduced by our model, but appear much more pronounced in the experiment.

### 3.2 Double heat shock

In [Schroda et al., 2000], a study was presented in which (for wider purposes than studying the HS response) an ARS gene, encoding for the enzyme arylsulfatase (ARS from now on) is placed under control of the HSP70A promoter. The authors show that whenever the HSP70A gene is activated also the ARS enzyme is produced. Under the assumption of a direct proportionality between the concentration of the ARS enzyme and its activity, the authors could monitor the activity of the HSP70A promoter by measuring the ARS activity. This construct was then used to systematically expose *C. reinhardtii* cells to two subsequent heat shocks to find out what is the minimum time one needs to wait after the first heat shock to observe a full HSR also in the second heat shock. It turned out that a waiting time of around 5 h is required to re-establish the capability to induce a full HSR.

To compare the experimental results to our model simulations, we extended the model accordingly, also including transcription and translation of the ARS enzyme (for details see Section B.2 of the Supplementary Material). In Fig. 5 we display the simulation results (left panel) and the corresponding experimental data (right panel) redrawn from Fig. 7b of [Schroda et al., 2000], where two heat shocks of 30 minutes duration were applied with the intervals of 2, 3, 4, and 5 hours, respectively (and also the curve for a single 30 minutes-long heat shock is shown). It can clearly be observed that both in simulation and experiment the second heat shock response increases in intensity with increasing length of the interval between the treatment, reaching its full activity after around 5 h. Again, the model results are in good quantitative agreement with the experimental data.

**Figure 5:**
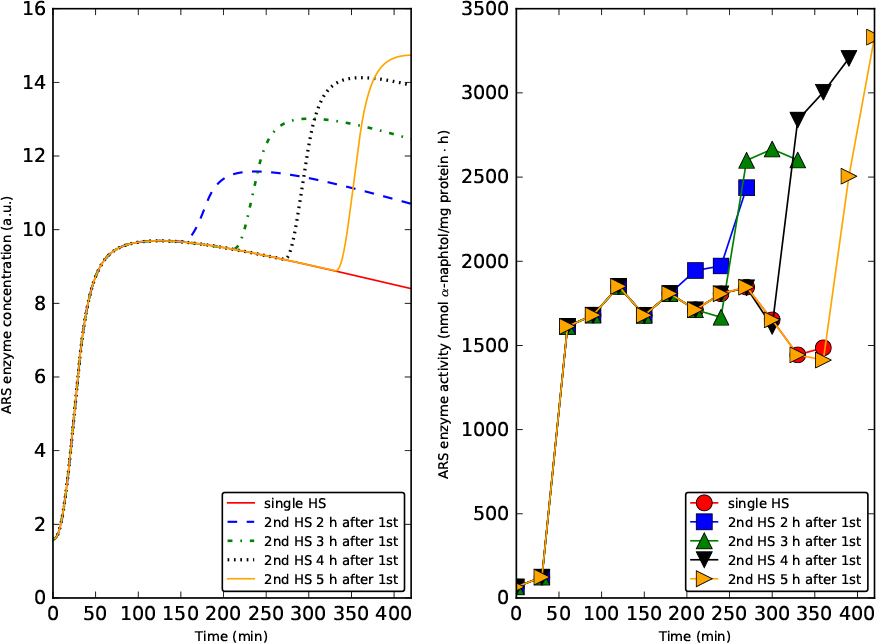
Simulating the double heat shock experiment (left) and comparison with the corresponding data from [Schroda et al., 2000] (right). We see that a full HSR is possible only about 5 h after the first HS.

An interesting observation when analysing the model simulations is that even after 5 hours, the concentration of HSP did not yet relax to its initial value before the first heat shock. While the second heat shock leads, as the first, to misfolded proteins and triggers an almost full HSR in terms of the observed HSP70 promoter activity, the amount of misfolded proteins is dramatically lower than during the first heat shock (see Fig. 12 and Section B.1 of the Supplementary Material). This indicates that the production of HSF resulting from the first HSR and the accumulation and slow degradation of HSP have the role of preparing the organism for a subsequent occurrence of the same stressing situation (the HS) already encountered in the past. Thus, the slow turnover of HSP may implement a short-term molecular memory of experienced heat stress, similar to the observed short-term memory of previously experienced light stress as was recently discussed and described by a mathematical model in [Matuszyńska et al., 2016]. Thus, the model simulations allows to give a novel interpretation of the experimental results on the double heat shock. While when observing the activity of the HSP70 genes seemingly a full heat shock response is observed, in fact the first and second response differ quite fundamentally. While during the first exposure HSP needs to be synthesised de novo from practically zero concentration and therefore the corrective response to refold the denatured proteins is slow, in the second heat shock after five hours the residual HSP level is sufficient to rapidly counteract the temperature-induced denaturing of proteins and the total level of misfolded proteins remains very low. However, even this low level is sufficient to induce expression of the HSP genes, so that on the mRNA level the second response appears as equally strong as the first.

### 3.3 Model calibration

The design of the model is based on our understanding of the underlying signalling network and is to a large part based on the findings reported in [Schmollinger et al., 2013]. As every model, our model is a simplified representation of reality. For example, a single variable *HP* describes the numerous HSP present in *C. reinhardtii*, and all genes involved in the HSR are described to possess the same promoter properties. While such a simplification may be seen as a weakness, because not all known factors are represented, the process of simplification itself is rather a very powerful tool, by which the essential features of the system can be extracted and it can be understood how the key properties of the dynamic response are generated. As a consequence, simplified models usually possess a more generic character, because the heat shock response may differ in details from organism to organism, but as long as the basic principles are conserved, our model serves as a generic theoretical description that can easily be adapted. Moreover, in a system such as the heat shock response investigated here, most model parameters are not directly accessible experimentally. In particular, the twenty rate constants listed in Table 2 of the Supplementary Material are largely unknown. This again supports the development of a simplified model, attempting to reduce the number of unknown model parameters as much as possible to facilitate a reasonable model calibration procedure, while maintaining the essential structural system properties.

To find plausible and realistic parameters, we initially considered biologically reasonable ranges for all rate constants and manually tuned these until we could well reproduce the qualitative behaviour of the experimental data. These parameters are referred to as the “fiducial parameter set” and are reported in the second column of Table 2 of the Supplementary Material. The fact that a very reasonable fit to experimental data could be achieved with a straightforward manual fit relies on the simplicity of the model structure and the concomitant fundamental understanding of the effects of the single parameters on the model behaviour.

We then used this fiducial parameter set as a starting point for a deeper investigation of the parameter space. We have divided the available experimental data described in the previous sections in two groups, one used to calibrate the model and the other to illustrate that the model is able to reproduce the qualitative behaviour observed in other experiments, as follows. The first group, used for calibration of the model, comprises the controls for the feeding experiments of [Schmollinger et al., 2013]. These correspond to six curves representing the time evolution of the concentration of mRNA coding for HSF1, and six curves for the mRNA coding for HSP90A, under heat shock and no inhibitor treatment.

We have then defined an objective function reflecting the quality of the fit by a simple root mean square of the deviations of the model simulations and the experimental data. We have first performed a Monte Carlo scan of the parameter space to gain insight into its structure and then a gradient search to find a set of parameters which locally optimizes the objective function. In this way we determined the “final parameter set”, which we use for all the simulations presented in this work, and which is reported in the third column of Table 2 of the Supplementary Material. A more detailed description of the calibration is provided in Section C of the Supplementary Material.

The second group contains data not used for model calibration, but for verification that the model is able to at least qualitatively predict the behaviour of this set. It contains the data on Staurosporine and Radicicol treatments discussed in Section 3.1 and those from the double HS experiment discussed in Section 3.2. Comparison of the simulation results using the final parameter set with the experimental data demonstrates that the model is able to reproduce the qualitative, and often the quantitative (relative, not absolute on which we do not have nformation) behaviour of the data extremely well (see Sections 3.1 and 3.2).

## 4 Results

As shown above, our mathematical model, which has been calibrated to control experiments only, can reasonably well reproduce drug treatments as well as the double heat shock experiments. The agreement of simulation results and experimental data therefore supports the notion that our current understanding of the heat shock response is basically correct. The mathematical model can therefore serve as a theoretical framework in which data can be interpreted in a sophisticated and quantitative way. Another purpose of model building is the ability to make novel predictions. We have therefore employed our model to simulate scenarios which give insight into our understanding of the heat shock response, but which are either difficult to test or have not yet been tested experimentally.

### 4.1 Prolonged heat shock

We first investigate which response the model predicts upon exposure to a prolonged HS, and how the system adapts to persistently high temperatures. Experimentally, the systems-wide response to long-term HS was investigated in [Hemme et al., 2014], where cells adapted to 25°C were exposed to 42°C for a period of 24 h, followed by 8 h at 25°C (recovery phase).

The simulation results for this scenario, where the temperature increase was simulated at time *t* = 0, are shown in Fig. 6. Two distinct phases of the response can be distinguished. The first, early HS phase, lasting approximately 3 h after applying the heat shock, represents the initial heat shock response, in which the internal variables respond highly similar to the normal heat shock simulations described above (see in particular Fig. 2, and also the controls curves in Fig. 3 and Fig. 4). During the late HS response, lasting from 3 to 24 h during heat shock, the variables approximate a new stationary state. This state is characterized in particular by increased concentrations of mRNAs (panel E), and consequently of HSF and HSP when compared to the corresponding concentrations before the onset of the HS. These elevated levels indicate an acclimation of *C. reinhardtii* to continuous HS conditions, which allows to efficiently avoid the accumulation of misfolded proteins. After reverting the conditions to normal temperatures (25°C), a recovery phase can be observed, in which the variables relax to the original stationary state over a period of several hours. These results are consistent with the observations of [Hemme et al., 2014] focusing on HSP production (see in particular Fig. 8 therein).

**Figure 6:**
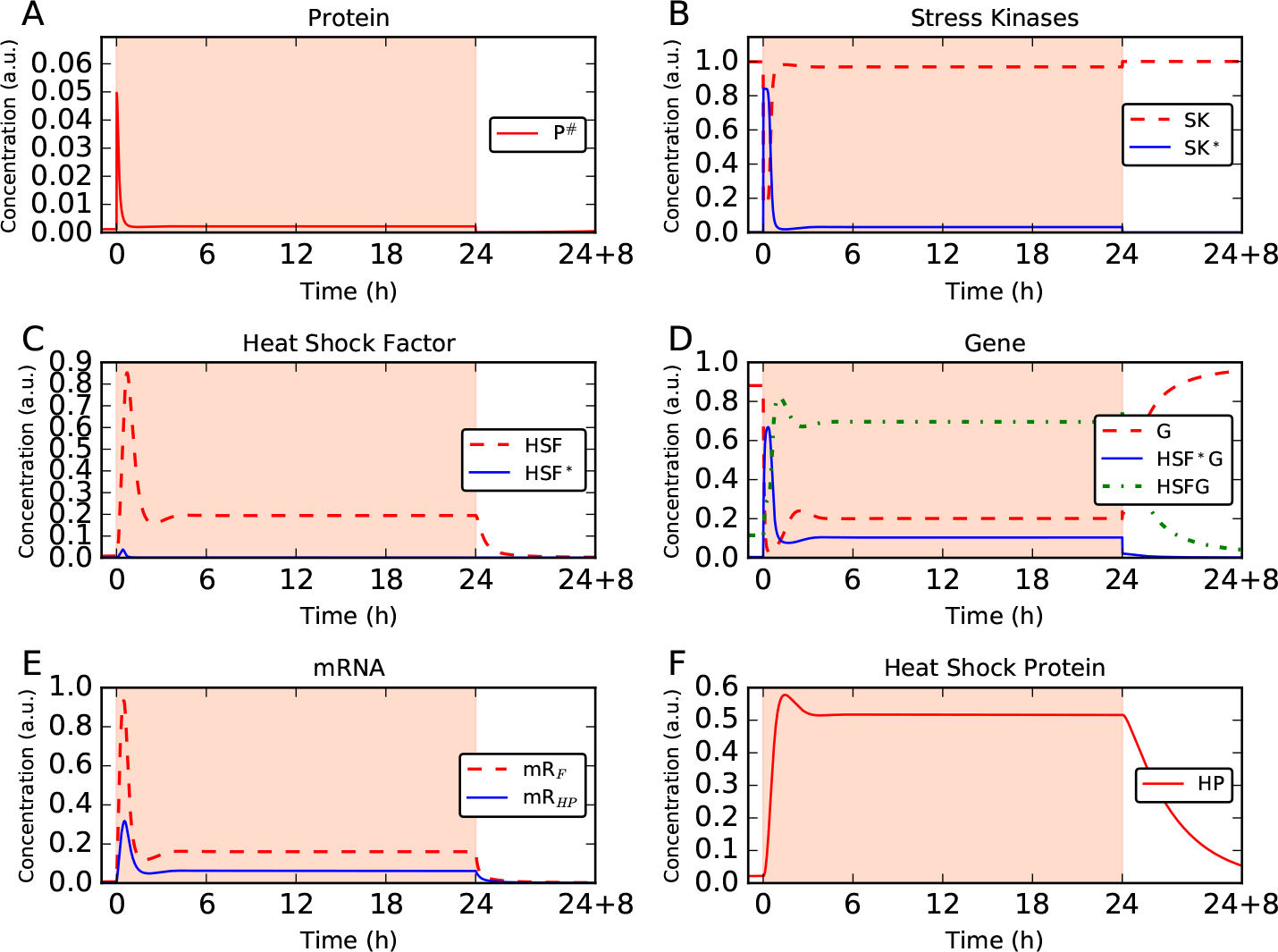
Simulation of the HSR under long-term HS and recovery, provided by shifting the temperature from 25°C to 42°C at *t = 0* and back to 25°C after 24 h. Two distinct phases are clearly visible: an early HS lasting for about the first 3 h, and a late HS in which the system shows adaptation (a new steady state is reached). After reverting the conditions to normal temperatures (25°C), a recovery phase can be observed, in which the variables relax to the original stationary state over a period of several hours.

### 4.2 Stationary behaviour

The observations that the system adopts a new stable stationary state when continuously exposed to elevated temperatures raises the question how this long-term response depends on the rate of protein denaturing. The steady state concentrations depend on temperature only via the speed of the reaction *P* → *P*^#^, which itself depends on the temperature, and on other quantities, as described by the corresponding equation in Table 4 of the Supplementary Material based on the Arrhenius law. We now study how the steady state concentrations depend on the temperature at which the steady state is reached.

First we have intuitively verified that the system reaches a steady state by naively integrating it over very long times. We then do it by applying the rigorous procedure which consists in looking for a root of the system represented by the ODEs of Table 3 of the Supplementary Material, i.e. to find the concentrations which correspond to a steady state. It’s best to have a good initial guess as a starting condition for the search, and the integration over a very long time performed above provides that. Then, we repeat this for different values of the temperature, to see how the concentrations change as a function of temperature. The result is plotted in Fig. 7. Let us remark that on the horizontal axes of Fig. 7 we have the temperature. Each point in a panel corresponds to the concentration of the corresponding species at steady state, when the steady state is reached at that particular temperature. We can see that the concentrations at steady state are different for different temperatures, with for instance higher values of the concentrations of mRNAs and HSP corresponding to higher values of the rate of the temperature.

**Figure 7:**
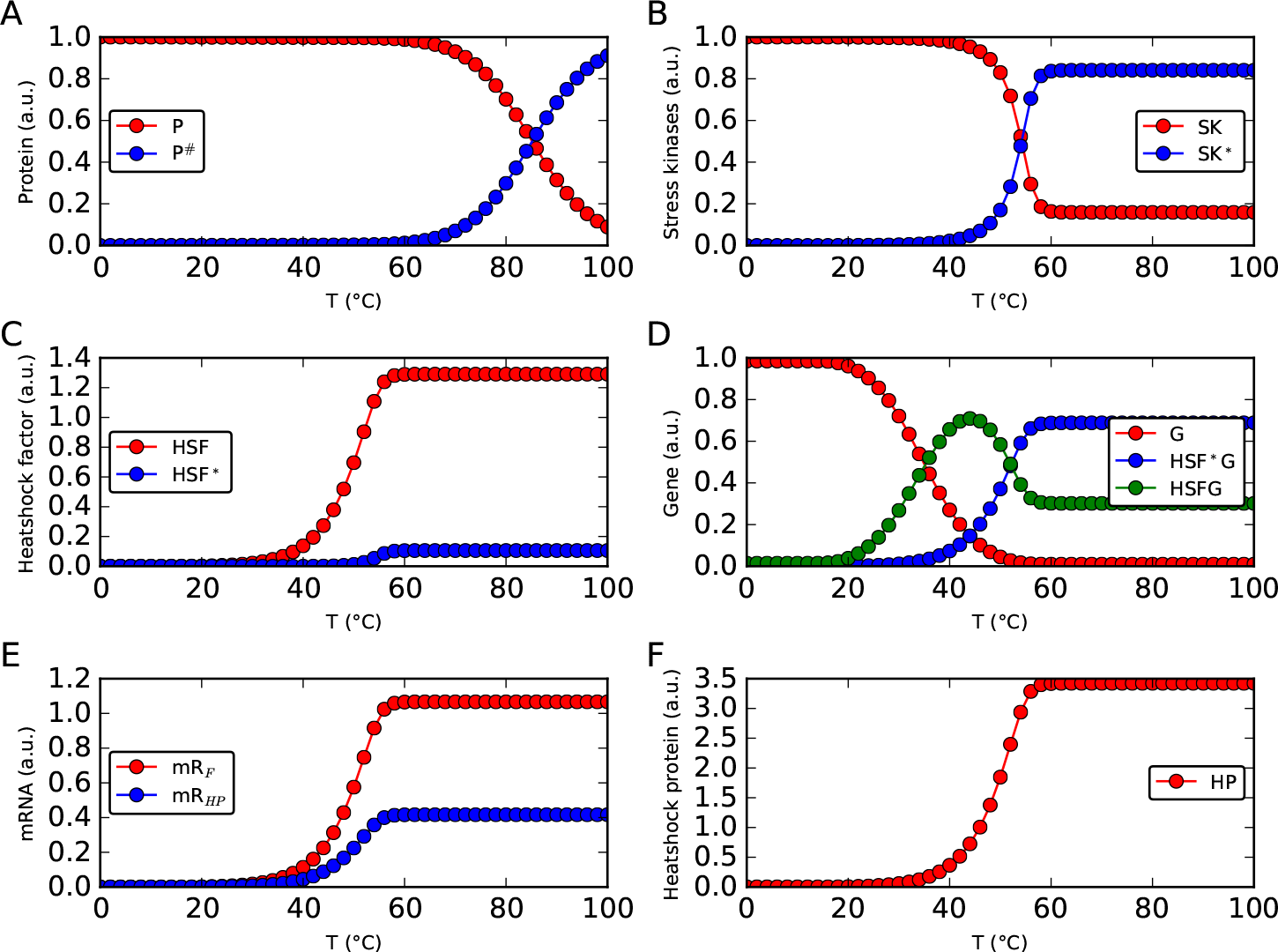
These simulations illustrates how different steady state concentrations are reached for different temperatures. Each point represents the value of the corresponding concentration at the steady state reached for that particular temperature. The concentrations of HP, as well as of the mRNAs, increase with increasing temperature. The concentration of unfolded proteins [*P*#] is kept very close to zero for low values of the temperature. When the temperature increases considerably the HSR is no more able to efficiently counteract the accumulation of degenerated proteins which accumulates at concentrations high enough to kill the cell. This accumulation is evident in panel A, a magnification of which is provided in Fig. 8.

On the one hand, for values of the temperature not too high, remarkably the concentration of unfolded proteins at steady state is kept very low by the response at the level of all the other species, very close to zero and in particular well below one percent of the total amount of proteins. On the other hand, when the temperature increases considerably the HSR is no more able to efficiently counteract the accumulation of degenerated proteins which accumulates at concentrations high enough to kill the cell. This accumulation is evident in panel A of Fig. 7, a magnification of which is provided in Fig. 8. Finally, we have also verified that, for each of the values of the temperature that we have considered, the model exhibits a realistic stationary behaviour, i.e. the associated steady state (which is a non-equilibrium one) is stable. To do so, firstly we computed numerically the Jacobian of the vector field associated to the ODEs system summarized in Table 3 of the Supplementary Material. Next, we evaluated the Jacobian of the system at the steady state under investigation. Then we computed the eigenvalues of the Jacobian to determine the stability of the steady state. We repeated the procedure for the steady state associated to each of the values of the temperature considered. We obtained that all the nine eigenvalues have always negative real part, showing that the steady state is always a stable one.

**Figure 8:**
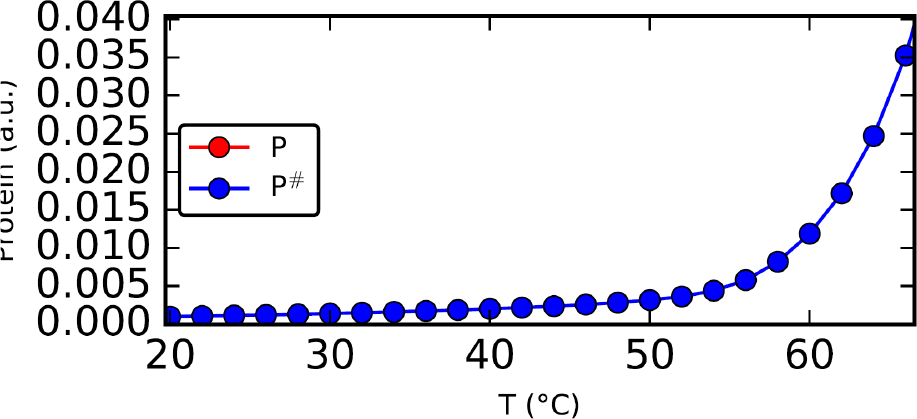
Magnification of panel A of Fig. 7, to emphasize the exponential growth of [*P*#] with increasing temperature.

### 4.3 Trade-off between HS temperature and HS duration in the production of HSP

The questions which we want to address in this section are the following: “is *C. reinhardtii* more stressed for a short HS with high temperature or a long one but with a lower temperature? What is the trade-off between the two?”. A related question is “does the production of HSP occurs only under very intense HS conditions (high temperature, long duration) or does it occurs also for very small temperature increases or very short HS?”.

Moreover, HS may also represent a mean to make proteins (HSPs, but not only) accumulate into plant cells. This might be interesting for instance in view of enriching plants in any protein of interest (by engineering the HSPs genes and use their HS-activable promoter to induce the expression of other genes of interest). The question that naturally arises is then: “which is the HS set-up (duration, temperature) that maximizes the accumulation of HSPs?”.

To answer these questions we performed a systematic study of how much HSP is produced under different combinations of HS durations and HS temperatures. For such a study, we provided to the system a sharp increase in temperature starting from 20°C. Since here the goal is mainly to study under which conditions *C. reinhardtii* is more stressed, and the response is closed only when HSP is produced and can act to refold unfolded/mis-folded proteins, and the main interest in eliciting an HSR may be to induce the accumulation of HSP (or other proteins), we perform this study at the level of HSP production.

Thus we simulate the response to the different combinations of HS temperature and HS duration, and we plot the value of [*HP*] computed right at the end of the HS period as contours in the plan representing HS duration versus HS temperature, and a colour map is used to make the figure visually clearer. It is worth to point out that this time point provides a value of [*HP*] that is not necessarily the highest one that can be obtained with an heat shock of that temperature and duration, in fact [*HP*] grows under HS, reaches a maximum, decreases a bit and settle to a new steady state value until HS is kept on. For short HS, as seen previously, even if the increase in temperature is sharp, and the activation of SK follows, there is a certain inertia in the response at the level of mRNA production, and an additional delay in the HP synthesis. As a consequence if the concentration of HP is read out at the end of a short HS, it is possible that the obtained value is lower than the value one would obtain with the same HS, but waiting some more time.

Fig. 9 shows the concentration of HSP at the end of the HS, as a function of HS duration and HS temperature. Firstly, looking at how [*HP*] changes for any fixed temperature, we can appreciate the same features that we already noticed in Fig. 2. There is no response at the level of [*HP*] for duration of less then about 10 min, than HP rapidly goes up for longer HS, and the maximum HP concentration occurs around 80 to 100 min. For longer HSs [*HP*] is somewhat lower, and does not change any more when increasing further the duration of the HS (new steady state reached, i.e. acclimation occurred). Second, considering increasing temperatures, we can see that even a small increase of few degrees in temperature results in an increase of [*HP*]. We can conceptually divide the plot in four regions (from left to right). No matter what temperature, for short HS (i.e. shorter than about 10 min) there is not enough time to lead to a significant increase in [*HP*]. For durations between approximately 10 min to 80 min and temperature above an increasing value, the contours are almost vertical. For high enough temperature, the HSR is activated and, no matter how much temperature is increased, the dynamics of the response is the same (this comes from the behaviour described by the Arrhenius law by which temperature enters the model by unfolding proteins). In the region of the maximum of [*HP*], the contour lines are almost horizontal, telling us that the maximum reached by [*HP*] strongly depends, and increases with, the temperature of the HS. Finally, on the right side of the plot we see that the contour lines are now horizontal and [*HP*] does not change with the duration any more: the system is acclimated (and thus has reached a new steady state). The values of [*HP*] are somewhat smaller than those corresponding to the region of the maxima.

**Figure 9:**
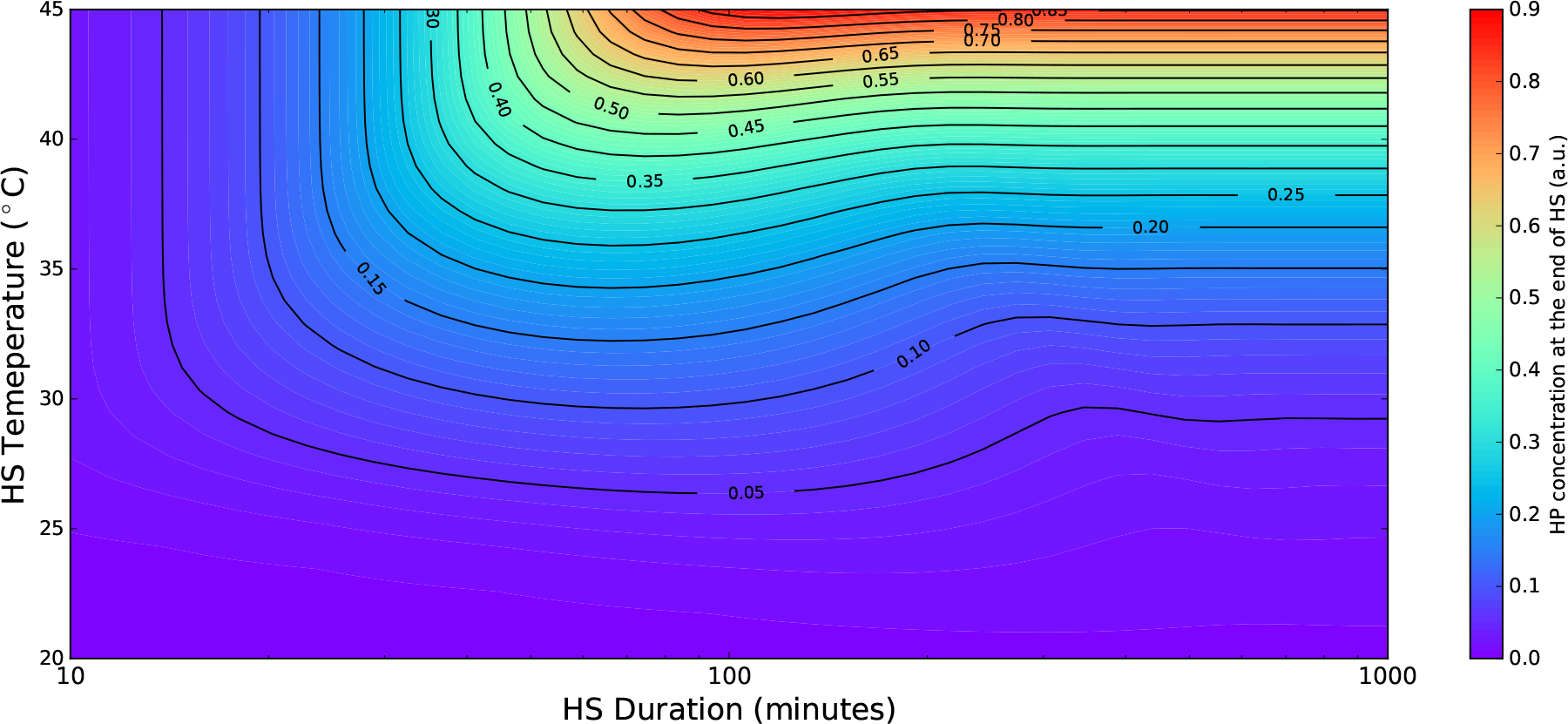
Systematic study of the HSP production as a function of different HS temperatures and HS durations. Short (smaller than 10 min) HSs do not provide enough time for a significant response at the level of [*HP*], the maximum of [*HP*] for any given temperature is obtained at around 80 to 100 min, after that a somewhat smaller [*HP*] is reached and maintained, and for long enough HS a higher temperature provides higher [*HP*]. From this plot one can understand the trade-off between duration and temperature.

### 4.4 The HSR to the temperature variation representing a hot day

To reproduce the typical experimental set-up employed in many studies we have considered throughout this work heat shocks provided by means of a step-wise increase of the temperature from a lower value to an upper value. These are conditions that can be easily reproduced experimentally, and allows to make straightforward tests in vivo. They could also be employed for possible applications of HS (as for instance expressing proteins of interest. Nevertheless, these are situations not so likely to occur often in nature.

Thus, which kind of HS is a *C. reinhardtii* cell going to experience in the wild? This green algae is widely distributed around the world in various environments such as soil and fresh water. Thus, a natural heat shocking condition it encounters is the daily variation of the temperature, which is low at night, grows during the day, reaches a peek and then drops again (with possibly some random fluctuations in addition). We then simulate an idealized variation in temperature reproducing that of a hot day, by imposing a sinusoidal variation between 22°C and 40°C with a period of 1 day and maximum at 3 p.m. (shown in Fig. 10), and we use our model to simulate the response of the system.

**Figure 10:**
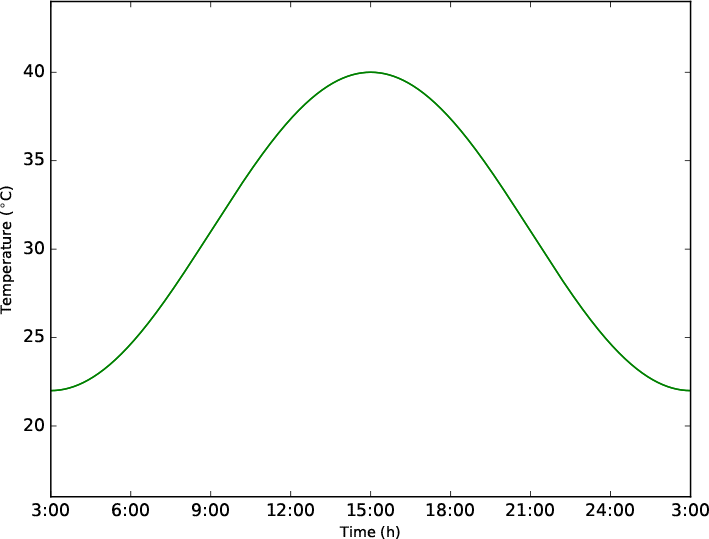
Temperature variation reproducing that of a hot day, used as input for the simulation of Fig. 11.

As we can see from Fig. 11, the concentrations of the mRNAs (panel E) and of the HSF (panel C) have a steep increase, which leads to a maximum and then a much slower decrease. The concentrations are not symmetric as the stimulus given to the system is. In particular in panel A we can appreciate the peak in the concentration of degenerated proteins at around 6 a.m.. This is due to the activation of the SK, which follows a Hill kinetics behaviour, which means that there is a threshold in [*P*#] above which SK become active and in turns activate the HSR. It is this HSR which lowers the concentration [*P*#] after 6 a.m. The response remains on until the evening and during this time the growth of [*P*#] due to the increase in temperature is counterbalanced by the HSR (this balance originates the second peak in panel A). When the temperature becomes sufficiently low in the evening, and [*P*#] as well, the HSR is switched off until the next day.

**Figure 11:**
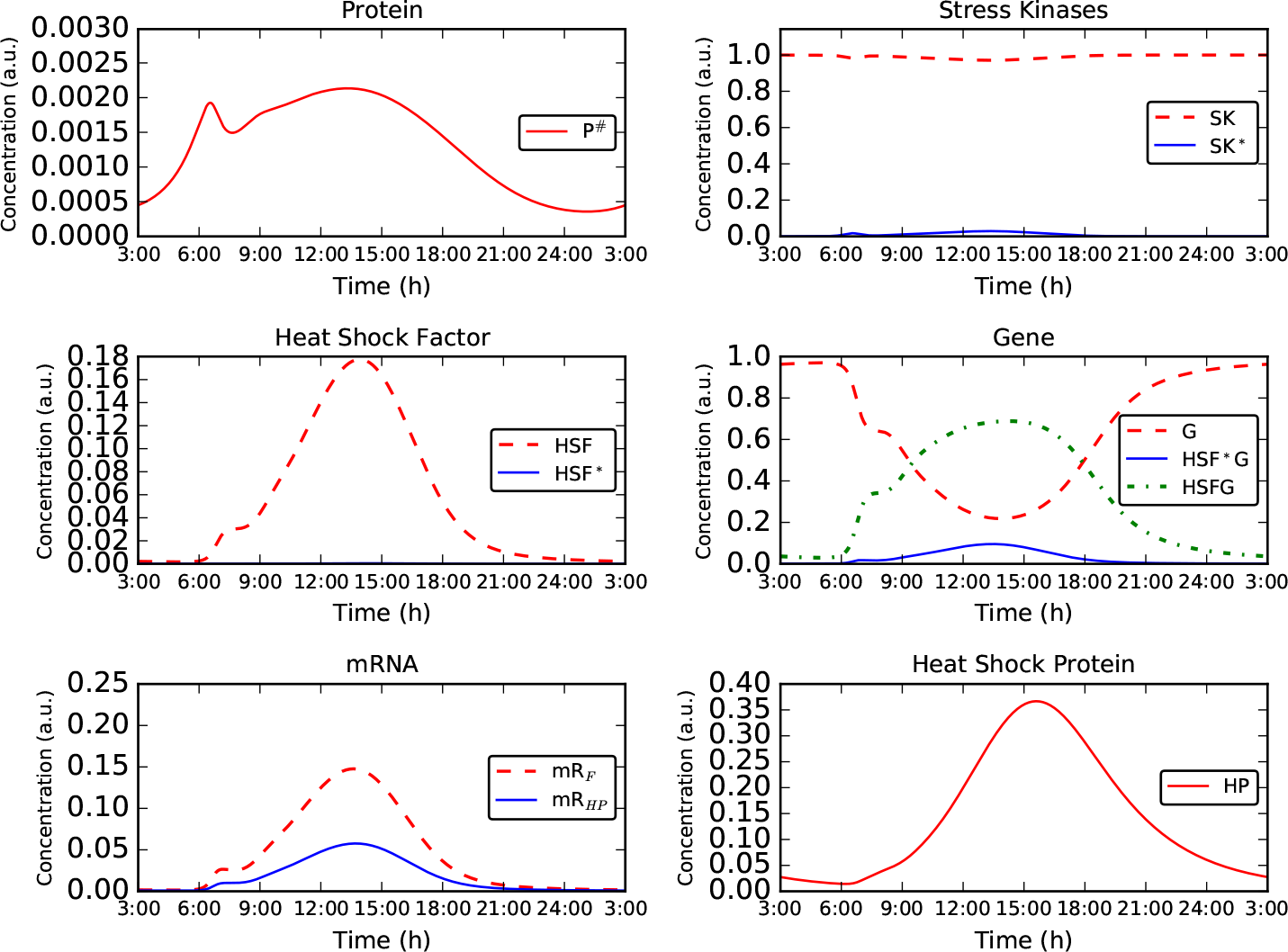
Simulation of the response of the system to a temperature variation reproducing that of a hot day (shown in Fig. 10). The concentration of unfolded proteins is kept very low, well below one percent of the total amount of proteins [*P*] + [*P*#] (panel A, where you can notice that the vertical scale is magnified w.r.t. the same panel of the previous figures).

Let us point out that the accumulation of unfolded proteins remains very close to zero (ways smaller than 1% of the total amount of proteins [*P*] + [*P*#], panel A) and in particular more than one order of magnitude smaller then when an HS of the same temperature is provided by a stepwise increase (panel A of Fig. 2). Since unfolded proteins are undesired by the cell, it is meaningful that the HSR, which for a sudden increase of 20°C is not fast enough and allows for a certain transient accumulation of unfolded proteins, is on the other hand perfectly capable to prevent the accumulation of unfolded proteins during a HS like those that can more often occur in nature. A systematic study on how the maximal concentration of HP changes when changing the time that is needed to increase the temperature from the ambient value to a higher value providing HS is presented in Section D of the Supplementary Material.

The Circadian clock of *C. reinhardtii* is well studied [Mittag et al., 2005, Jacobshagen et al., 2001], it is known to regulate also some HSP as e.g. HSP70B, and it provides to the concentration of this protein a periodic behaviour with maximum at dawn. The HSR that we model is not a Circadian clock, but we see that it gets activated slowly just before dawn, rapidly goes up to a maximum before midday (some hours before the maximum of the temperature), drops slowly for the rest of the day, is roughly off when night comes, and stays off over night. It seems that the cell, as soon as an increase in temperature is sensed, starts an HSR to prepare itself to the higher temperature that is going to come, and then during all the day it “keeps the defences up” to deal with the increased temperature, and it turns them off only when the value of the temperature is sufficiently dropped. All this by keeping always the concentration of unfolded proteins close to zero.

## 5 Conclusions and Outlook

In this work, we have built a data driven mathematical model for the HSR in *Chlamydomonas reinhartii*, a photosynthetic model organism. We have introduce the model in Section 2. We have described the signalling pathway underlying the model, we have presented the mathematical description of the mechanism that we implement, we have discussed the typical behaviour of the model and we have verified that the system is stable.

We have extracted the signalling network structure from various experiments, and experimental data are used for calibration and validation of the model. The comparison with experimental results extracted from the literature is described in Section 3. We have first described which experimental data we consider (feeding experiments and double HS experiments), we have then discussed how we use them for calibration of the model and we have finally used them to show that the model is able to reproduce experimental data. The capability of this model to reproduce the main qualitative features of various experimental datasets shows that our conclusions about the signalling mechanism are plausible. Moreover, the model allows for analysing the response on different signal levels which are not easily accessible in experiments.

In Section 4 we have used the model to simulate situations not yet tested experimentally, from which we derive interesting results. We have shown in Section 4.1 that the system can adapt to higher temperatures during heat shocks longer than three hours, by shifting to a new steady state. Two distinct phases are clearly visible: an early HS lasting for about the first 3 h, and a late HS in which the system shows adaptation (a new steady state is reached). The recovery phase is characterized by a recovery of the conditions pre-HS that occurs over several hours.

We have studied in Section 4.2 the variation of the steady state concentrations w.r.t. changes in the temperature. The concentrations of HP, as well as of the mRNAs, increase with increasing temperature, but for not too high temp)eratures the concentration of unfolded proteins [*P*#] is kept very close to zero, in particular well below 1% of the total amount of proteins [P] + [*P*#]. For temperatures too high the HSR cannot prevent the accumulation of degenerated proteins and the cell dies.

We have used the model to systematically investigate how the accumulation of HSPs depends on the combination of temperature and duration of the heat shock, in Section 4.3. Short (smaller than 10 min) HSs do not provide enough time for a significant response at the level of [*HP*], the maximum of [*HP*] for any given temperature is obtained at around 80 to 100 min, after that a somewhat smaller [*HP*] is reached and maintained, and for long enough HS a higher temperature provides higher [*HP*].

We have finally investigated the system response to a smooth variation in temperature simulating a hot day in Section 4.4, showing that the percentage of proteins which are unfolded remains well below one percent of the total amount of proteins.

It is then natural to wonder what would be interesting to investigate next using this model as a starting point. A number of observations (listed below) shows that the HSR could be elicited in *C. reinhartii* also by a shift from dark to light of the cells, by an independent regulatory pathway. We thus propose that our model could be extended in the future to include and investigate how this occurs. This extension could help investigating the potential role of an intermediate of Chlorophyll biosynthesis, Mg-Protoporphyrin IX, as a mediator in the signalling pathway between the chloroplast and the nucleus.

It is known from the eighties that some of the genes coding for HSPs in *C. reinhartii* are inducible by a shift of the cells from dark to light (as shown in [von Gromoff et al., 1989] which experimentally proved that for genes of the three families HSP68, HSP70 and HSP80). The two regulatory pathways leading to the activation of these genes by a shift from dark to light on the one hand, and by HS on the other, have been shown to be independent in [Kropat et al., 1995] in the case of the HSP70 family of genes. Therein it is also shown that the kinetics of the two pathways is different, and that an additional protein synthesis should be involved in the light-activation pathway.

To understand better how this could work, we need to understand the structure of the HSP genes, for instance the HSP70A gene, and in particular of its promoter region, which are well described for instance in [Lodha and Schroda, 2005] and [Strenkert et al., 2013]. On the promoter region of this gene, four sites are present where HSF1 can bind and activate one of the two transcription sites leading to the transcription of the gene itself.

In [von Gromoff et al., 2006] a detailed study is carried out with the conclusions that on the proximal region of the above mentioned promoter there are two additional binding sites, which would allow for another molecule, namely Mg-Protoporphyrin IX (Mg-Proto), to activate one of the two transcription sites present on this promoter, thus activating the transcription of the gene. [Kropat et al., 2016] studied in more detail the role of the chloroplast signalling in the light induction of nuclear HSP70 genes, which (they conclude) requires the accumulation of chlorophyll precursors and their accessibility to cytoplasm/nucleus. In [von Gromoff et al., 2008] a mechanism is proposed to explain this: Mg-Proto is synthesized in the chloroplast, but when cells are kept in the dark it cannot exit this organelle. When cells are shifted from dark to light, some channels are opened which allow Mg-Proto to exit the chloro-plast, become available in the cytoplasm and reach the nucleus, where they can bind to the promoter region of the HSP70a gene, thus activating its transcription. [von Gromoff et al., 2008] also shows that a similar role can be played by Heme, whenever Mg-Proto is less available.

Moreover, with a certain degree of speculation, the positive feedback loop involving a further protein synthesis might potentially be closed in the following way. It is known that one of the very first steps of the biosythesis of chlorophyll (and thus of Mg-Proto, an intermediate) requires the action of an enzyme called Glu-Tr. This enzyme is coded for by a gene called HEMA. [Vasileuskaya et al., 2005] has shown that Mg-Proto and Heme control the expression of this HEMA gene. This could represent the positive feedback loop which would allow the full activation mechanism to be speed up when light would make Mg-Proto (or Heme) available outside the chloroplast. Moreover, a negative feedback loop is also known to be present in the biosynthesis of chlorophyll, namely if too much of some products of Mg-Proto accumulates into the chloroplast, the very first steps of this biosynthesis are inhibited.

An extension of our model to include all these effects would thus present an interplay between two feedback mechanisms. On the one hand, the positive feedback on the production of Mg-Proto due to Mg-Proto itself made available (by light) outside the chloroplast, which can activate the HEMA gene, necessary for the very first steps of Chlorophyll biosynthesis. On the other hand, the negative feedback on the production of Mg-Proto due to Mg-Proto itself accumulating inside the chloroplast. Developing this possible extension of our model, while besides the scope of this work, would provide a theoretical and quantitative framework to investigate further the potential role of the intermediate of Chlorophyll biosynthesis Mg-Protoporphyrin IX as a mediator in the signalling pathway between the chloroplast and the nucleus.

Moreover, the HSP genes are involved in helping the cell to face other types of stress. For instance their role in helping the recovery from photo-inhibition is investigated in [Schroda et al., 1999], and evidence for protection by heat-shock proteins against photo-inhibition during heat-shock is discussed in [Schuster et al., 2016]. These represent further interesting directions in which our model could possibly be extended.

## Download of our code

The code by which we have implemented the model and the simulations hereby described can be freely downloaded from the following git repository: https://github.com/QTB-HHU/ModelHeatShock.

## Acknowledgements

We are grateful to all the members of the Institute for theoretical and quantitative biology, Düsseldorf, for very useful discussions about this work, and to Simon Schliesky for the informatics support that made it possible. This work was financially supported by the European Union Marie Curie ITN AccliPhot (PITN-GA-2012-316427) to S.M., A.S. and O.E., and Deutsche Forschungsgemeinschaft Cluster of Excellence on Plant Sciences, CEPLAS (EXC 1028) to O.E.

## Author Contributions

Conceivement and design of the study: OE, ASku. Code development: SM, ASku. Model development: OE, ASku, SM. Analysis and interpretation of the results: SM, ASuc, ASku, OE. Drafting the manuscript: SM, ASuc, ASku, OE. Coordination of the study: OE.

## A Supplementary Material: mathematical description of the model

## B Supplementary Material: further study of the double heat shock

In this section we present some additional simulations which illustrate extensively the features of the response to a double heat shock, and next we explain how we have extended our model to simulate the double HS experiments performed in [Schroda et al., 2000].

### B.1 Further simulations of double HS experiments to illustrate their main features

We present in Fig. 12 a series of four figures which illustrate how the HSR dynamics changes when we simulate a generic double heat shock experiment, providing to the system two HSs at a distance of 30 min, 2 h, 3 h 30 min and 5 h respectively. For this simulation we employ a sudden variation of the temperature between 25°C and 42°C.

**Figure 12:**
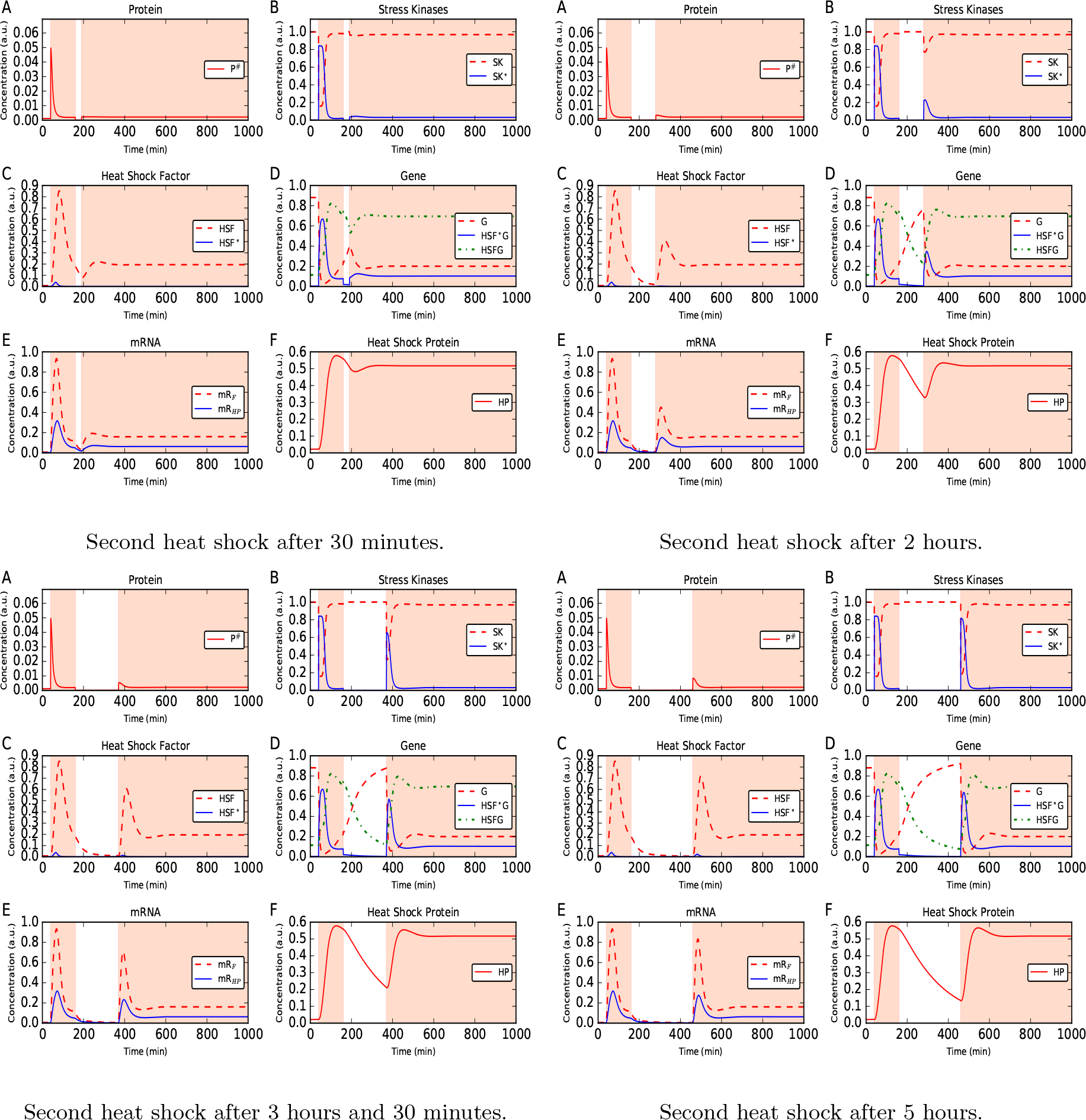
Simulation of a generic double heat shock experiment. The second heat shock is provided respectively after 30 min, 2 h, 3.5 h and 5 h. We can appreciate how the dynamics changes at the level of each species. Particularly relevant is the fact that a full response to the second HS is possible only after about 5 h, as clearly shown by e.g. the SK*, HSF* and mRNAs curves.

As we can see from Fig. 12, when the second HS occurs only 30 min after the first, the system shows almost no response to the second HS. This is due to the fact that during the first HS, thanks to the gene activation (panel D) and subsequent production of mRNA for the HSF (panel E), the quantity of HSF available to the system increases (panel C). When the second HS occurs, 30 min after the end of the first, the HSF available to the cell is already enough and no activation of the SK takes place (panel B). When the second HS occurs a lot of HSP is still available in the system (panel F) and thus the level of degenerated protein *P*# does not increase (panel A).

When the second HS occurs 2 h after the first, there is a small HSR during the second HS, that we can see at the level of the SK (panel B) and of the mRNAs (panel E). This because even if a lot of HSP is still available (panel F), the HSF concentration is very low (panel C) and then a moderate HSR is necessary to allow the system to quickly refold unfolded proteins. The HSR corresponding to a second HS occurring at 3 h 30 min after the first is similar, but enhanced. When the second HS occurs 5 h or more after the first we see that the concentration of HSP is approaching the level it had before the HS (panel F), all the other quantities are approximately back to the original values, and an almost full HSR takes now place when the second HS is applied (panels B, C, D and E).

It is very interesting to remark that, while the concentrations of all the species go back to the values that they had before the first HS quite fast after the end of the first HS, the HSP does not (panel F), and this allows to avoid during the second HS having any but a tiny amount of unfolded protein *P*# with respect to the amount during the first HS. This can be seen in panel A of any of the sub-figures of Fig. 12, where the concentration of degenerated proteins *P*# increases by a considerable amount during the first HS, while considerably less during the second even when this is occurring many hours after the first.

The behaviour we observe in our simulations likely indicates that the production of HSF which follows a first HSR and the accumulation and slow degradation of HSP have the role of preparing the system for a subsequent occurrence of the same stressing situation (HS) already encountered in the past, thus representing a transient molecular memory.

This behaviour is qualitatively consistent with the claim of [Schroda et al., 2000] that *C. reinhardtii* needs around 5 h to recover and be able for another full HS response. We have seen in the main text a direct comparison of simulations performed with our model and data from [Schroda et al., 2000], and in the next section we provide additional details on how we have extended our model to perform such simulations.

### B.2 Extending the model to simulate the experiments performed in [Schroda et al., 2000]

To perform the simulation reproducing the data of [Schroda et al., 2000], shown in Fig. 5 of the main text, we have extended our model including few new variables and equations. We included the production of ARS enzyme into our model by adding two new variables and four new reactions to it. The two variables represent the concentration of the mRNA coding for the ARS enzyme, due to fusion of the ARS gene on the HSP70A promoter, and the concentration of the ARS enzyme itself. The four new reactions that we add describe the production of mRNA coding for the ARS enzyme from G*HSF, the translation of ARS mRNA into ARS enzyme (i.e. production of ARS), a very slow degradation of the ARS enzyme and degradation of ARS mRNA.

We selected the values of the free parameters in order to match the observations. In Fig. 13 we studied the behaviour of the added part of the system and we compare it with experimental data, verifying that the qualitative behaviour is similar.

**Figure 13:**
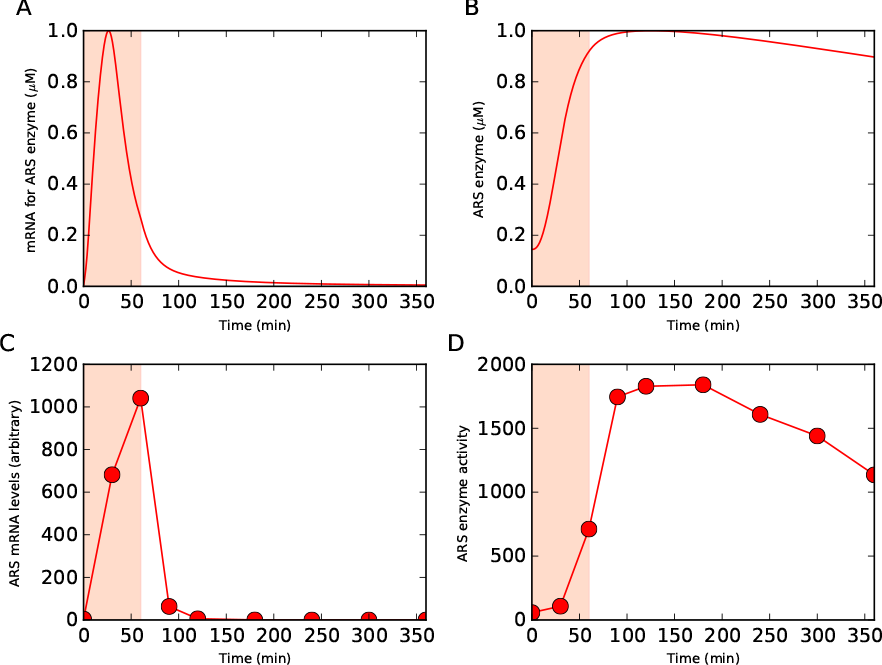
Comparison of simulations (upper panels) with data from [Schroda et al., 2000] (lower panels) on single heat shock experiment to verify that the extension of our model allows to reproduce the behaviour observed therein.

Applying an HS of 1h by increasing the temperature from 23°C to 40°C we have an increase of the concentration of the mRNA for the ARS enzyme due to the increase of HSF*G in our model, which reaches a peek and then drops (panel A). We also have an increase of the concentration of the ARS enzyme delayed w.r.t. that of the mRNA (due to the additional time necessary for protein synthesis), and which has a much slower attenuation (panel B) so that it takes ways longer to the ARS concentration to drop that to the mRNA concentration. We can compare the predicted concentration of mRNA with the measured one (panel C, data from [Schroda et al., 2000], and the predicted concentration of ARS enzyme with the measured ARS activity (panel D, data from [Schroda et al., 2000], Fig. 6b). This comparison requires the simplifying, but still reasonable, additional assumption that the activity of the ARS enzyme is roughly directly proportional to its concentration. Given the simplicity of the extension that we used to include this into our model, and the rough estimation of the involved parameters, the qualitative agreement between simulation and data is already remarkable.

## C Supplementary Material: extended description of the calibration of the model

In this section we provide more details on the procedure that we have employed to calibrate the model. We describe in detail the objective function used and which data are involved in calibration. We then describe the random sampling of the parameter space and the gradient search employed to determine the final set of parameter values.

### Definition of the objective function

To assess how well the model simulations reproduce the behaviour of the corresponding data points from [Schmollinger et al., 2013], we employ as objective function the root mean square deviation (RMS) of the theoretical prediction w.r.t. the data on mRNA expression for HSF and HSP, for the six controls of the feeding experiments (which we called previously the first group of data). It is important to point out the, as can be seen from the figures of [Schmollinger et al., 2013], the data, obtained with northern blot analysis, correspond to relative concentrations and not to absolute concentrations, i.e. the data points of each control curve are normalized to the maximum among all the time points and all the curves (controls and feedings) for that feeding experiment. Thus, to compute the RMS we normalize the values of the concentrations to the maximum of each curve, to be able to compare these with the experimental data. For this reason we can only say that the model can reproduce the qualitative and the quantitative (but relative, not absolute) behaviour of the system: the data employed do not contain absolute (dimension full) measures of the concentrations, but only relative measures (i.e. normalized to a maximum). Thus, while the qualitative and quantitative (relative) behaviour of the simulations performed with the model are calibrated on the data and thus reliable, the absolute quantities which it provides represent only a reasonable indication of the possible values assumed by the concentrations, and for this reason they appear in each figure normalized to a reference value (see Table 5).

#### C.1 Random exploration of the parameter space employing a Monte-Carlo analysis

The fiducial parameter set from which we start, manually tuned by means of our understanding of the mechanism underlying the model, already allows the model to roughly reproduce the qualitative behaviour observed in the experiments. Nevertheless, we decided to explore more systematically the parameter space represented by the twenty rate constants of Table 2, and study how the RMS just defined changes if we move around from the fiducial parameter set.

For this purpose we first performed a Monte Carlo (MC) scan of the parameter space. We did so by assuming a flat prior probability distribution between half and two times of the fiducial value of each parameter. The part of the parameter space which we explored is thus a 20-dimensional hypercube, every point having the same probability of being randomly selected. Then, by randomly extracting a value for each parameter from these distributions, we generated 10^5^ randomized sets of parameters.

For each randomized set of parameters we computed the corresponding value of the RMS with respect to the data of the first group, used for calibration. We obtained values of the RMS ranging approximately from 0.13 to 0.70. The fiducial parameter set has a RMS w.r.t. the controls of the feeding experiments of 0.147, which is already remarkably close to the lower edge of the range obtained using random combinations of the parameters.

We then select the 5% of the points corresponding to the best (i.e. lowest) values of the RMS (which then range between 0.130 and 0.149). By plotting for these points the value of the RMS versus the value of each parameter (an example is shown in Fig. 14), we observe that for the vast majority of the parameters no preferred interval in which the lowest values of the RMS occur more often can be identified. Very low values of the RMS can occur everywhere in the interval used for the random scan, depending on the values assumed by the other parameters.

**Figure 14:**
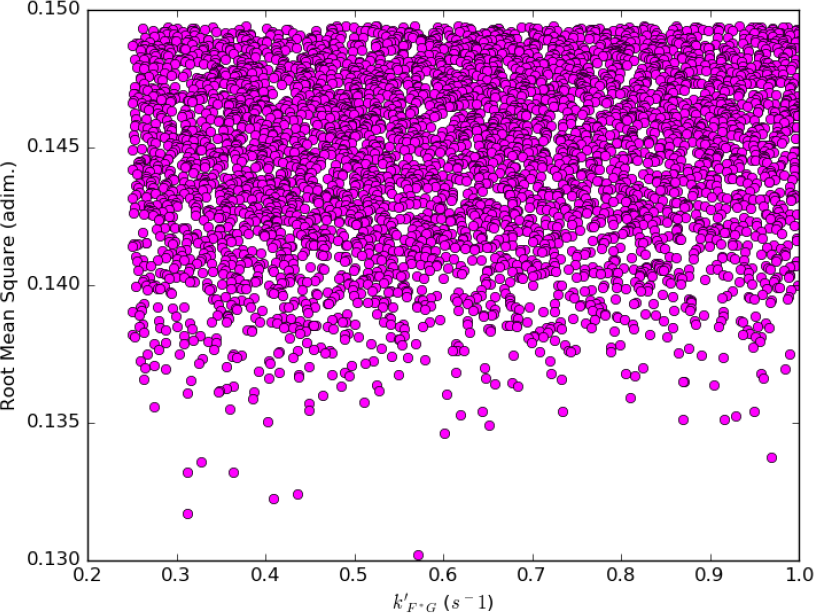
Example showing RMS vs 1 of the model’s parameters for the points corresponding to the 5000 random parameter sets with lowest RMS. We repeated this analysis for the 20 parameters, obtaining similar results.

We subsequently plotted the values of the RMS as function of each couple of parameters (obtaining 400 figures). We found that sometimes there is some correlation or anti-correlations between the preferred values for the two parameters of the couple (an example is shown in Fig. 15). Nevertheless, for the majority of the couples no such correlation can be identified.

**Figure 15:**
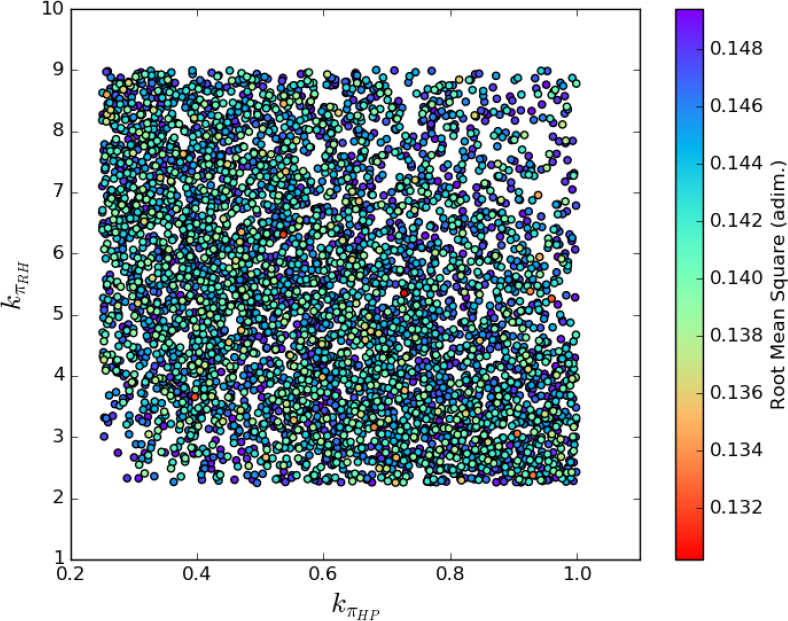
Example of figure showing RMS (colour coded) as function of two of the model’s parameters for the same points of Fig. 14. We repeated this analysis for the 400 combinations of two model’s parameters at a time. This one is among the few showing (anti) correlation.

Moreover the parameter set which provides the best RMS among this randomly generated set (i.e. the isolated point at the bottom of Fig. 14) turns out to be not a good one, because if used to run the simulations of other situations as for instance the double HS, it provides to the model a behaviour qualitative completely different from what is observed in the experiments.

These observations lead us to conclude that many very different configurations (distributed almost everywhere) in the parameter space would allow us to obtain a small RMS with respect to the data, but in any case this small RMS would not be that much smaller than the value corresponding to the fiducial parameter set.

On the other hand, if we consider small variations of one parameter at a time, we observe smooth variations in the RMS values, showing that the RMS looks roughly parabolic for perturbations of each one of the majority of the parameters around the corresponding fiducial value, and that the fiducial value often lies close to the minimum (as shown in the example of Fig. 16).

**Figure 16:**
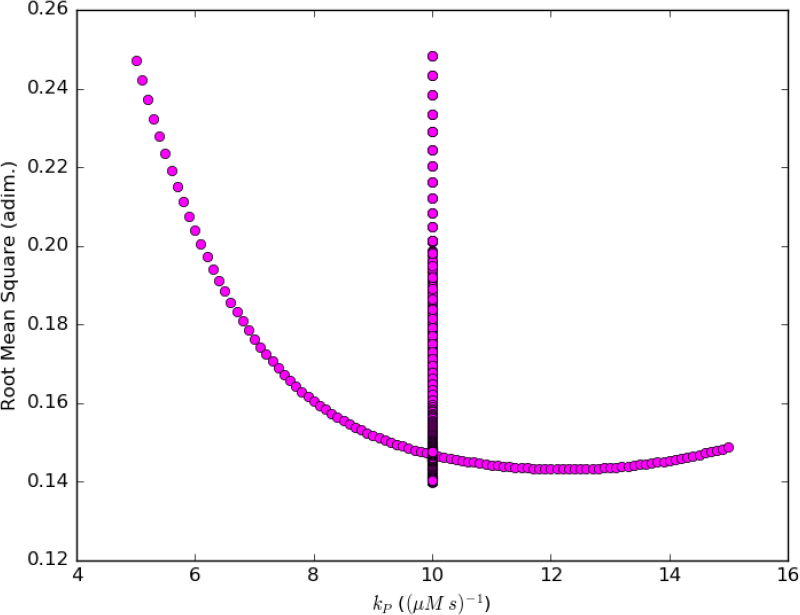
Example of figure showing RMS versus one of the model’s parameters, when we consider variations only in one of the parameters at a time. The points aligned along the vertical line in the center of the figure correspond to variations in one of the other parameters, and are showed for comparison. We repeated this analysis for the 20 parameters of the model.

Thus, after having performed a global random scan of the parameter space to gain a better understanding of how the RMS behave in it, we decided to determine the final parameter set by employing a local optimization method, namely a gradient search starting from the fiducial parameter set.

This provides a parameter set allowing a better fit to the data used to compute the RMS, while not moving too far away from our already well behaving fiducial set. It is important here to recall the generality of the model’s description of the HSR, and the lack of measurements on the parameters involved. Thus the goal of the model is, given an input (the temperature as function of time), to simulate a plausible output (the behaviour of the concentrations as functions of time), and not to determine the “internal” parameters involved (the rate constants).

As a final test we defined, similarly to what we have done above, a RMS distance between the model simulations and the experimental data from the double HS experiment of [Schroda et al., 2000] (reported in Fig. 5). Defining such an RMS is much more arbitrary than defining the RMS w.r.t. the controls of the feeding experiments, because of the previously mentioned hypothesis on the proportionality between the enzyme concentration and its activity. We do not combine the two RMSs, as this would require to attribute to the two a weight which would be highly arbitrary.

For these reasons we only compute this second RMS a posteriori as a check, for the best 5000 among the random parameters sets selected above, and show the distribution of the values of the two different RMS in Fig. 17. This shows that minimizing both at the same time cannot be obtained, and one needs to find a trade-off between the two. The values of the RMS w.r.t. double HS run between 0.11 and 0.55 (not all visible in the figure, which is magnified). We can a posteriori compute this RMS for the fiducial parameter set, finding 0.170, and for the final parameter set, obtaining 0.168.

**Figure 17:**
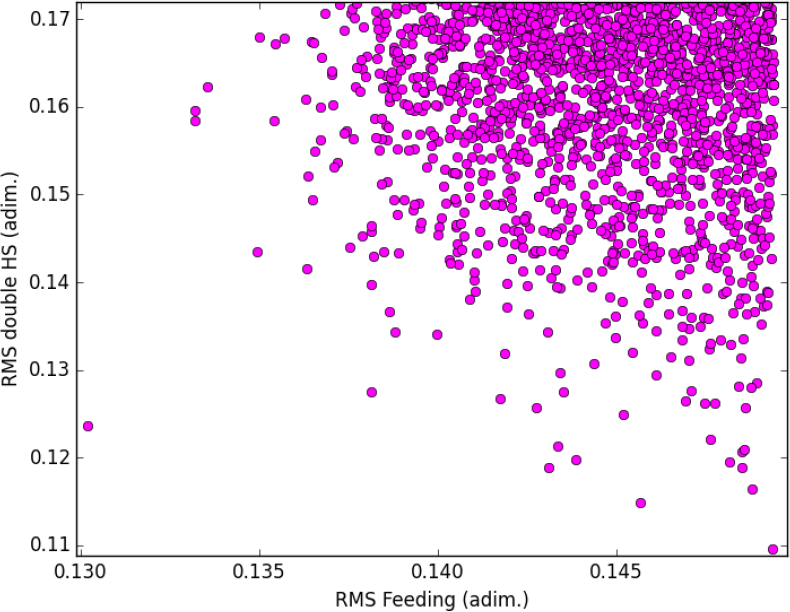
RMS w.r.t. the double HS data versus RMS w.r.t. the controls of the feeding experiments, magnifying the region where the points with lowest values of the RMSs lie.

Finally, we verified that in the gradient search performed in the next section, if we employ the sum of the two RMS instead of the RMS w.r.t. to feedings only, we can improve the fit to the double HS data, but at the cost of having non-realistic behaviours in the model consisting of big oscillations in the concentrations after onset of HS (and of introducing an arbitrary weight between the two RMSs).

#### C.2 Local optimization using a gradient search to fit the parameters’ values to the data

As discussed above, the fiducial set of parameters allows already to obtain a value of the RMS (w.r.t. the controls of the feeding experiments) close to the lower limit of the random set. We explored the parameter space with the MC analysis and we concluded that the RMS can be improved only marginally w.r.t. the value corresponding to this set. Having shown that no region of the parameter space is preferred by the RMS, we opted for a local optimization procedure, thus performing a gradient search starting from the point represented by the fiducial set of parameters, employing the steepest descent method.

This means that we start from a point 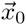 in the parameter space represented by the fiducial value for the 20 rate constants employed in our model (see Table 2). We compute the corresponding value of the root mean square *RMS* 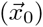. We then compute numerically the gradient of the RMS at that point 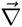*RMS* 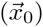 (by approximating partial derivatives using the symmetric difference quotient).

We then proceed along the direction opposite to the gradient towards a new point 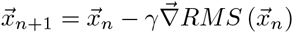 in the parameter space which provides a smaller value of the RMS. We do so iteratively until a termination criterion described above is satisfied, and label the iteration number by *n*. At each iteration, we need to decide which is the length of the step *γ* that we want to use in the direction opposite to the gradient. To do so, we implement a line search with the aim to loosely minimize the function 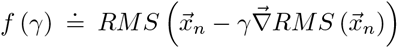 w.r.t. *γ*, i.e. along the direction opposite to the gradient. This means finding the value of *γ* which minimizes the function *f* (*γ*). We do so numerically employing a modification of the bisection rule based on the Golden ratio to save computation time.

Since the orders of magnitude of the parameters are very different, we expect the isosurfaces of the function *RMS* 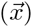 to be far away from being spherical. This would lead to a very slow convergence of the method, because the gradient at each step would hardly point roughly toward the minimum. We thus employed also a preconditioning of the function that we want to minimize, i.e. the RMS. We applied the numerical procedure of minimization not to the actual *RMS* 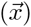, but to a function *RMS'* 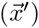 which we obtain by transforming *RMS* 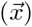 via a rescaling of all the parameters using their fiducial values. In this way all the parameters are of order of magnitude one, which is more suitable for the application of the described numeric algorithm. Once the minimization has been performed, we applied the inverse transformation to re-obtain the parameters in their original form.

The termination criterion we employed imposes the algorithm to stop when the average RMS decrease over the last ten iterations is lower then a threshold value. This criterion is of course somewhat arbitrary, but we have selected it by empirically verifying that it allows a better fit to the control data (a lower RMS), avoiding an over-fitting which would lead to model behaviour too far away from the behaviour expected w.r.t. other situations as e.g. the double HS, as can be seen from Fig. 18 and Fig. 19. The algorithm stopped after 31 iterations and returned the set of parameters listed in the second column of Table 2 as final values. The corresponding value of the RMS w.r.t. the controls of the feeding experiments is 0.137. We have employed this set of parameters to perform all the simulations shown in this work (a part from the calibration runs).

**Figure 18:**
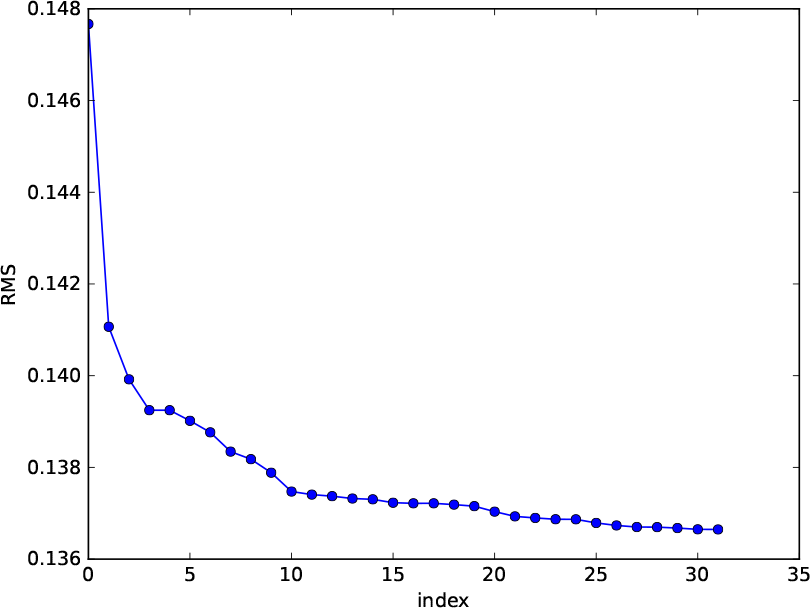
RMS decrease for subsequent iterations of the gradient search algorithm.

**Figure 19:**
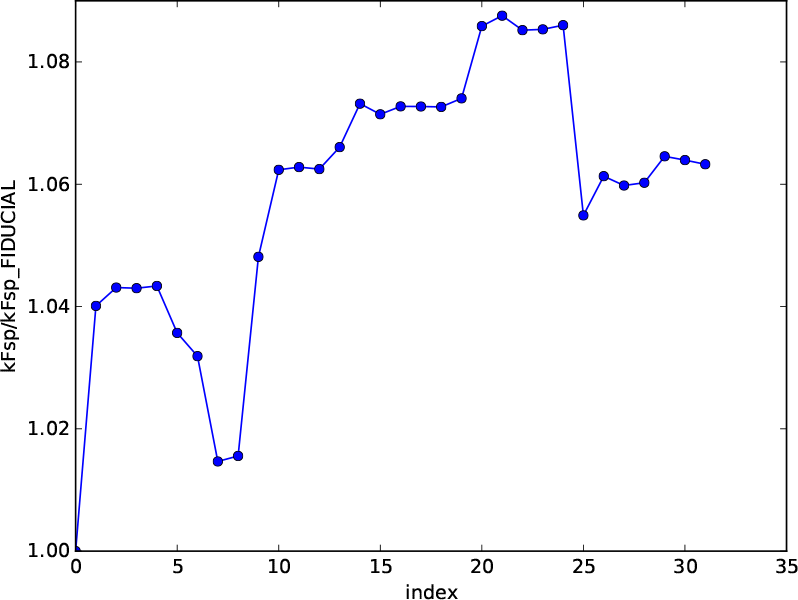
Values of one of the parameters for subsequent iterations of the gradient search algorithm.

#### C.3 The model reproduces the qualitative behaviour observed in the experiments

We have compared the results of our simulations to the corresponding experimental data, those of the second group. These are the double heat shock experiments performed in [Schroda et al., 2000] and shown in Fig. 12 and the feeding with Staurosporine and Radicicol performed in [Schmollinger et al., 2013] and shown in Fig. 3 and Fig. 4. Additional comparison with data on the expression of the HP protein collected in [Mühlhaus et al., 2011] are shown in Fig. 21 and discussed in the Supplementary Material.

The figures mentioned above show the corresponding results of the simulations performed with our model using the final parameter set determined as described in the previous section. The experimental set-ups, the salient features of each experiment and the way to use our model to simulate these experiments have been widely described in Sections 3.1 and 3.2. As we can see from the figures, the model reproduces well the qualitative and the (relative) quantitative behaviour of these experimental data.

## D Supplementary Material: Maximal accumulation of degenerated proteins as function of time necessary to increase the temperature during HS

We might wander how the maximum concentration of unfolded proteins accumulated during a HS depends on how fast the temperature has increased from the initial value *T*_*low*_, to the final value *T*_*high*_. To study this we simulate what happens when providing to the system a HS by acclimating the system at the temperature *T*_*low*_ =25°C, then starting from *t = t*_0_, then increasing the temperature following a cosinusoidal function until when the temperature *T*_*high*_ =42°C is reached at *t* = *t*_0_ + *τ*, and then keeping the temperature at *T*_*high*_. In this way the function *T* (*t*) is not only continuous, but also everywhere differentiable. We repeat for various values of the time *τ* which is the time required for the temperature to increase from *T*_*low*_ to *T*_*high*_, and we show the result in Fig. 20. We observe that for instantaneous increase in temperature, up to increases which requires about 1 minute, the maximum value of the concentration of degenerated proteins does not change. For *τ* between about 1 minute and 100 minutes there is a steep fall, and another plateau is reached for *τ* bigger then about 100 minutes. Notice that the scale on the horizontal axis of Fig. 20 is logarithmic.

**Figure 20:**
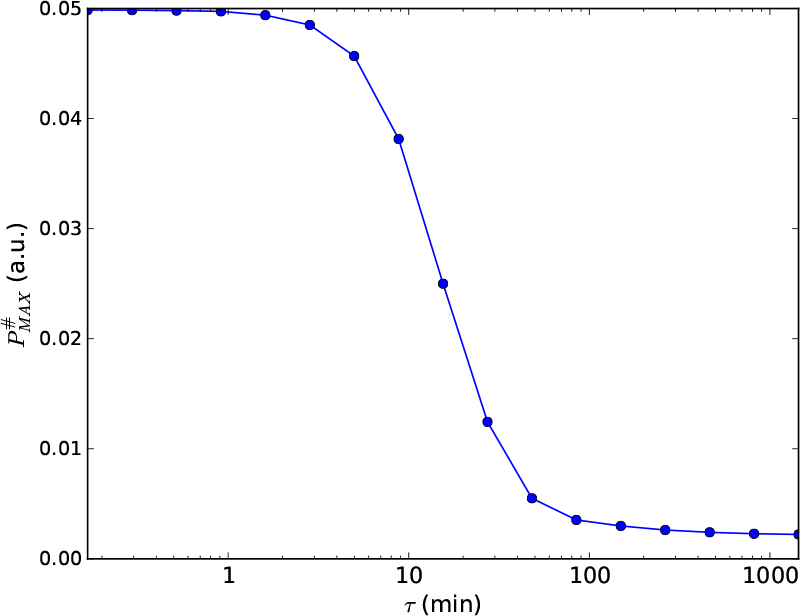
Study of how the maximum concentration of unfolded proteins accumulated during a HS depends on how fast the temperature has increased from the initial (lower) value to the final (higher) value.

## E Supplementary Material: comparison with data on HP expression from [Mühlhaus et al., 2011]

Since the comparisons among model’s predictions and experimental data worked out so far are mainly at the level of mRNA production, here we would like to compare with data the model’s predictions for the time evolution of the concentration of HP. We consider the data from [Mühlhaus et al., 2011], which using quantitative shotgun proteomics monitors proteome dynamics in time course experiments on *C. reinhardtii* and, among the thousands of proteins analysed, found 38 proteins significantly increased upon HS. To simulate this we apply at t=0 min an HS by increasing (as in the experiments) the temperature from 25°C to 45°C. We compared the model simulated behaviour of [*HP*] with the data relative to HSP70A, HSP70B and HSP90.

Fig. 21 shows that the model reproduces the qualitative behaviour of the data: the concentration of HP increases slowly at the very beginning, then faster, reaching a maximum and mildly decreasing afterwards. Quantitatively, the data are provided in terms of the z-score, a quantity used in statistic to describe a set of data points, which measures the distance of the single data point from the mean, in terms of standard deviations (thus a z-score of -1 means 1 standard deviation below the average of the set of data points). While not having access to the absolute values, we nevertheless know given its definition that the z-score is linear with respect to the concentration value represented by each data point, then we do not expect any distortion on the vertical axes of the plot if we would be able to transfer these data to the corresponding original values, which justify the comparison of Fig. 21. The qualitative agreement is good, nevertheless we can notice a somewhat faster increase during the first 100 minutes of the model’s prediction w.r.t. the data.

**Figure 21:**
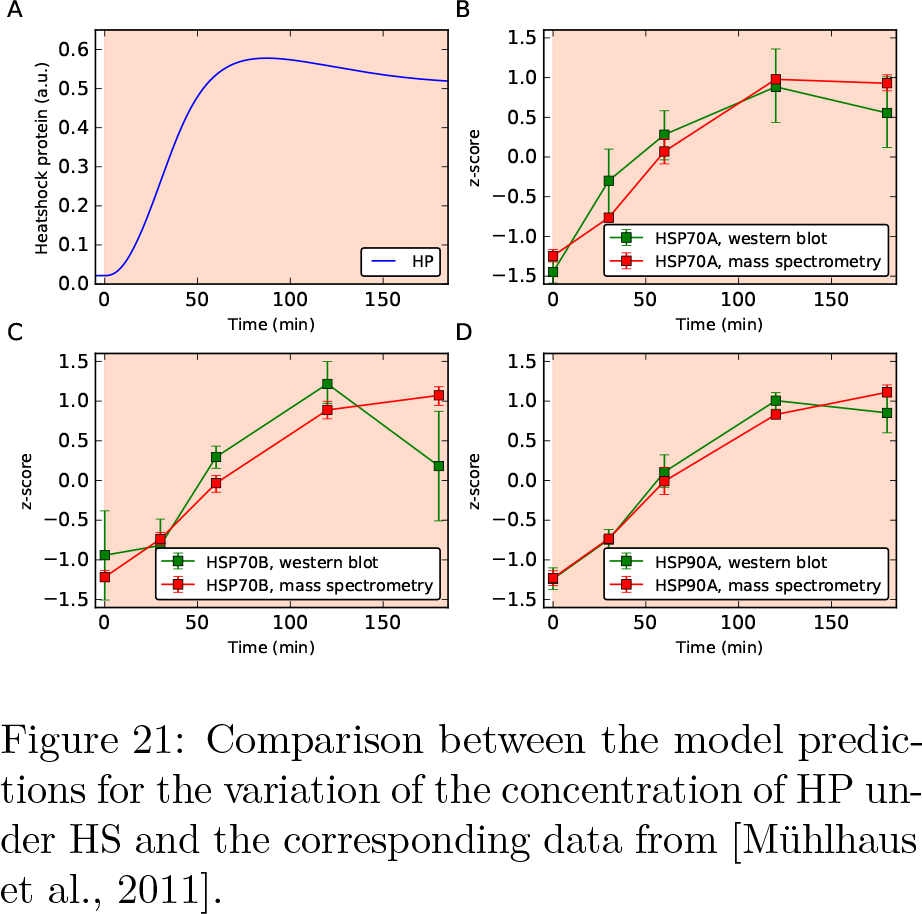
Comparison between the model predictions for the variation of the concentration of HP under HS and the corresponding data from [Mühlhaus et al., 2011].

